# Cell Line Development for Bispecific Antibodies: Better Predictability Through Transposases

**DOI:** 10.1101/2025.08.12.669435

**Authors:** Sowmya Rajendran, Indira Kottayil, Lynn Webster, Divya Vavilala, Molly Hunter, Monica Konar, Surya Karunakaran, Mario Pereira, Jeffrey Johnson, Jeremy Minshull, Ferenc Boldog

**Author notes:** These authors contributed equally to the work (Sowmya Rajendran, Indira Kottayil, Lynn Webster).

## Abstract

Bispecific antibodies are at the forefront of biopharmaceutical drug development. With over 100 different molecular architectures combined with diverse individual subunit sequences, choosing the most suitable structure and predicting the ideal subunit expression ratios for successful heterodimerization is a significant challenge. In this paper, we demonstrate that the recently described cell line development paradigm shift (Rajendran *et al*. 2021), enabled by the Leap-In transposon platform, can be extended to the development of bispecific monoclonal antibody-producing cell substrates (stable clones and pools). The key features are 1) Parental pools reliably predict the derivative clonal productivity and clonal heterodimer fractions. 2) Clonal productivity and clonal heterodimer fraction remained stable for at least 60 population doublings. 3) Depending on the products’ biophysicochemical properties, the stable pools exhibit variable productivity stability. 4) Heterodimer fractions remain stable in the Leap-In mediated stable pools independently of the productivity stability of the pools. 5) Structures and subunit ratios can be triaged at stable pool level, and 6) Due to the homogeneous clonal productivity distribution, only a small number (∼50) of clones need to be isolated and characterized.

## Introduction

One of the latest significant advancements in biopharmaceutical drug development is the successful application of multi-specific, predominantly bispecific, biologics. From a therapeutic perspective, a relatively few distinct classes of biologics can be identified (e.g., modulating two or more targets to achieve synergistic effects on a biological pathway, or redirecting effectors to specific targets with an enhanced therapeutic index (Deshaies, 2020); however, structurally there is a myriad of different molecular architectures designed to achieve these therapeutic objectives Spiess *et al*., 2015, Brinkmann *et al*., 2017, Godar *et al*., 2018).

Most bispecific architectures require heterodimer formation based on several immunoglobulin heavy chain pairing and heavy chain-light chain pairing designs (Madsen *et al*., 2024; Spiess *et al*., 2015. The manufacturing of these complex designs by mammalian cells still poses unique challenges. Novel protein expression technologies, including expression construct design and stable cell line development workflows, are needed to maximize heterodimer formation.

The recently introduced Leap-In transposase® system is well suited to address these challenges. (Rajendran *et al*., 2021). First, the structural and functional integrity of the stably integrated transposon-based constructs faithfully maintains the designed subunit expression levels at every integration site in every recombinant cell. Second, due to their efficiency and integration site selection, the Leap-In transposase mediated stable pools expressing single chain molecules and homodimers, like mAbs or Fc fusions, present a high level of productivity and product quality comparability commensurate with derivative stable clones. This unique feature enables the evaluation of various chain ratios on stable pool expression levels, identification of pools with optimal product quality attributes, and a limited single cloning effort for the isolation of clones with attributes reflected by the original pool.

A critical component of the Leap-In system is the transposon-based expression construct design. Since transfection efficiency presents the only limitation for integrating the transposon-based expression constructs, positioning of multiple ORFs with selected regulatory elements within one Leap-In construct is straightforward.

This manuscript demonstrates the robust stable pool to stable clone productivity and heterodimer fraction comparability, even without optimizing construct designs for maximum productivity and heterodimer formation. Furthermore, using both three-chain and four-chain bispecific antibody models. Stable pools and respective clones exhibit comparable productivity and heterodimer fractions while maintaining volumetric productivity and product quality stability.

## Materials & Methods

### Gene synthesis, vector construction

Emicizumab is a three-chain asymmetric bispecific antibody composed of one anti-FIXa heavy chain, one anti-FX heavy chain, and one common light chain. (Kitazava T., Shima, M. 2020) Figure 1a. The four-chain Vanucizumab is an example of the 1+1 CH1LC CrossMab architecture, with two different heavy chains and two different light chains targeting Angiopoietin 2 and Vascular endothelial growth factor A (Schaefer *et al*. 2011), Figure 1b.

**Figure 1.**
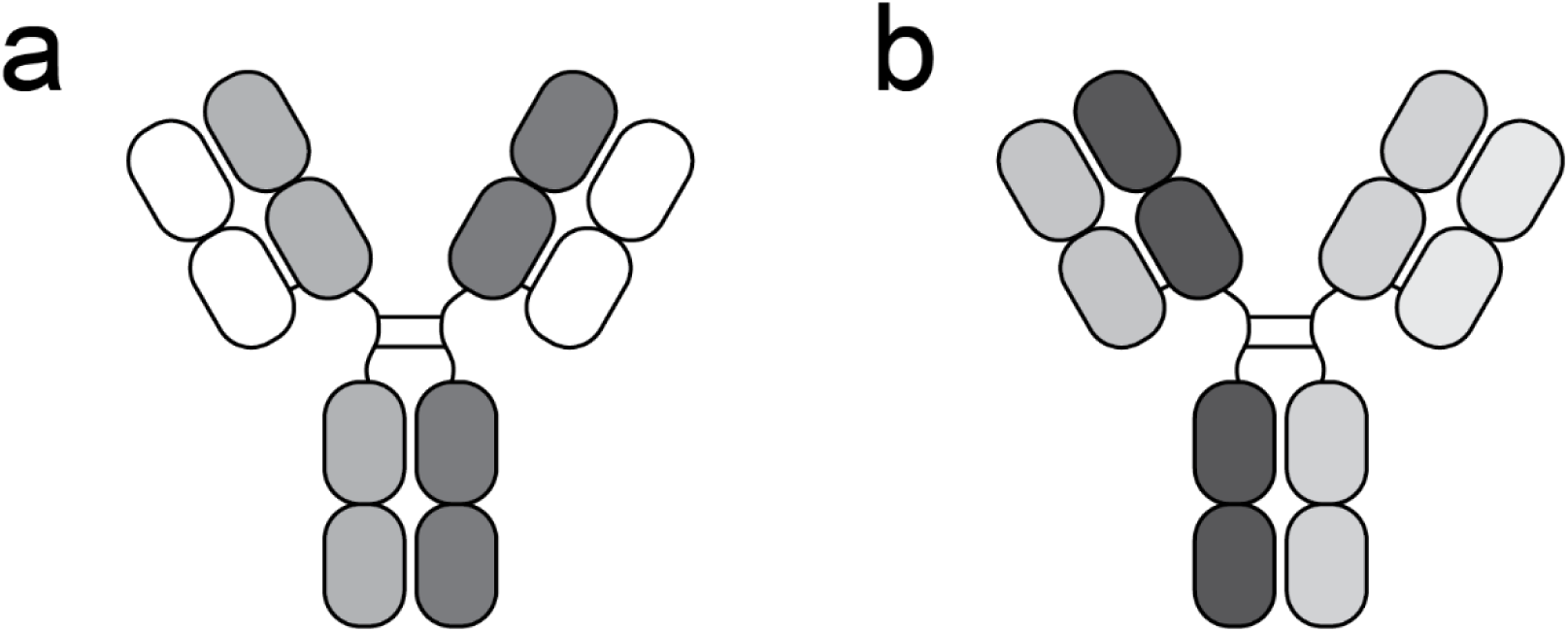
Schematic representation of molecules. The structure of Emicizumab (a). The molecule contains two different heavy chains and a common light chain—the structure of Vanucizumab (b), an example for the 1+1 CH1LC CrossMab architecture. The molecule contains two different heavy chains and two different light chains.

Schematic drawings of the Emicizumab and the Vanucizumab architectures are presented in Figure 1a and 1b, respectively.

The recombinant genes were synthesized and, together with the fully synthetic gene expression regulating elements, were assembled into transposon-based expression constructs in ATUM’s laboratories using proprietary technologies. The sequences of the assembled constructs were confirmed using Sanger or Oxford Nanopore Technologies sequencing methods.

Brief description of the expression constructs used in the study:

a. Ten Leap-In transposon-based expression plasmids, all coding for the three different Emicizumab chains at various expression ratios, were designed to establish ten corresponding stable pools presented in Table 1. The expression constructs represent combinations of Position1 5’UTRs, and polyadenylation sequences with Position2 5’UTRs and a common Position2 polyadenylation sequence. Different positioning of the transcriptionally linked LC-HC1 and LC-HC2 sequences represented another variable.
b. Nine Leap-In transposon-based expression constructs, all coding for the four different Vanucizumab chains at various expression ratios, were designed to establish nine corresponding stable pools Table 2, upper panel. These 4ORF constructs, conceptually similar to the 3ORF Emicizumab plasmids, represent a combination of different 5’ and 3’ UTRs for the LC1-HC1 transcript in Position 1. The LC2-HC2 transcript, in Position 2, was designed with different 5’UTRs and with a common polyadenylation site.
c. Two Leap-In constructs, expressing either the LC1 and HC1 or the LC2 and HC2 ORFs, have been designed for co-transfection at three different plasmid ratios. The LC1/HC1 and LC2/HC2 constructs were designed with the same 5’ and 3’UTRs with the LC sequences in the first and the HC sequences in the second positions. The corresponding pools are presented in Table 2. Lower panel
d. To support the development of analytical methods determining heterodimer content, constructs coding for the homodimers were also designed using the same 5’ and 3’ UTRs as in the multi-ORF constructs.
e. Constructs coding for the functional Emicizumab and Vanucizumab subunits carried a glutamine synthetase (GS) selection cassette, Leap-In transposon ends, two insulator sequences flanking the integrated transposons, as well as a kanamycin resistance gene and an *E. coli* replication origin. To minimize the deviation from the intended plasmid ratios in Vanucizumab pools 11, 14, and 15 established by co-transfection, the two constructs were built with the same GS selection cassette.

**Table 1.**
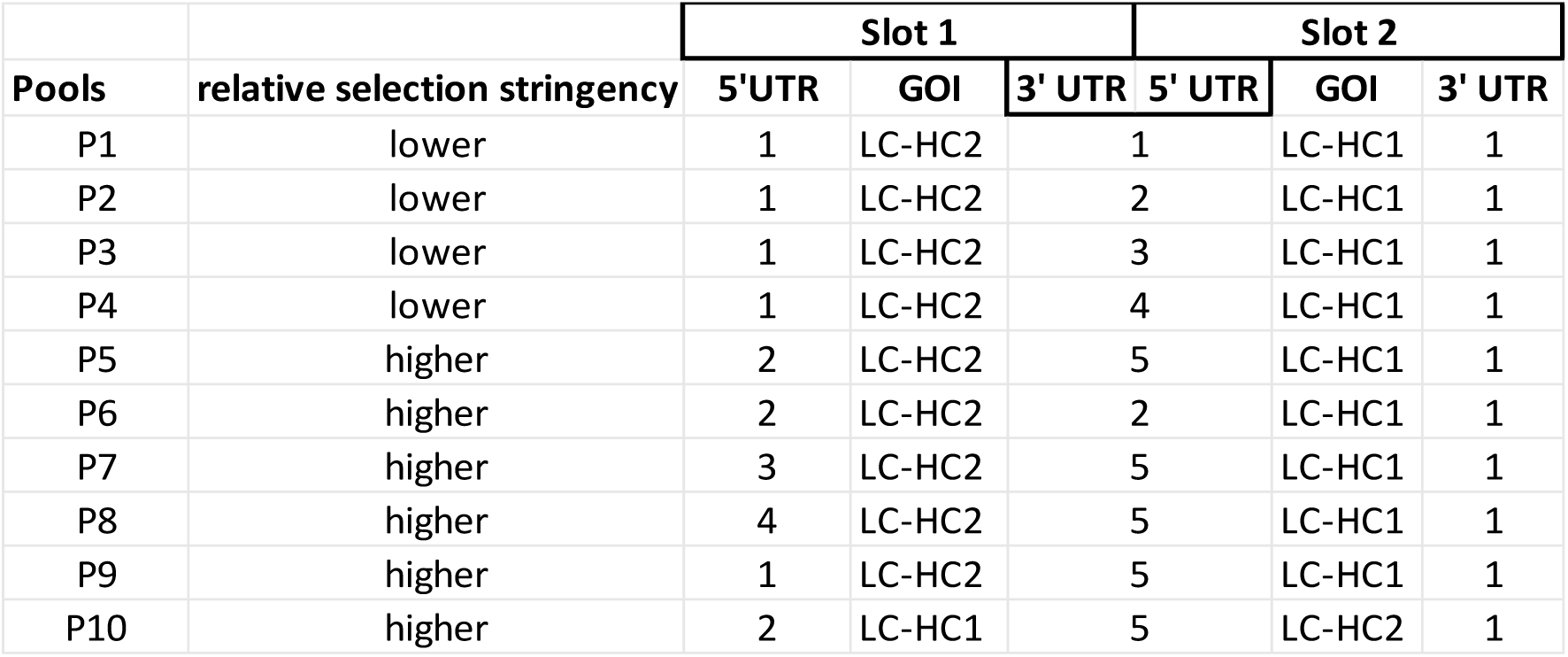
Regulatory element combinations in the Emicizumab expressing Leap-In transposon-based expression constructs. The numerals represent various regulatory element combinations. For example, five different combinations of Slot1 3’ UTR and Slot2 5’ UTR were designed.

**Table 2.** Regulatory element combinations in the Vanucizumab expression constructs. Upper panel: 4 ORF constructs; lower panel: 2 x 2ORF co-transfections. The numerals represent the different 5’ and 3’ UTR sequences.

| Pool | 5' UTR1 | 1<sup>st</sup> slot GOI | 3' UTR1 | 5' UTR2 | 2<sup>nd</sup> slot GOI | 3' UTR2 |
| --- | --- | --- | --- | --- | --- | --- |
| 2 | 1 | LC1-HC1 | 1 | 6 | LC2-HC2 | 4 |
| 3 | 2 | LC1-HC1 | 1 | 7 | LC2-HC2 | 4 |
| 4 | 3 | LC1-HC1 | 3 | 7 | LC2-HC2 | 4 |
| 5 | 3 | LC1-HC1 | 1 | 7 | LC2-HC2 | 4 |
| 6 | 3 | LC1-HC1 | 2 | 8 | LC2-HC2 | 4 |
| 7 | 4 | LC1-HC1 | 1 | 6 | LC2-HC2 | 4 |
| 8 | 5 | LC1-HC1 | 1 | 6 | LC2-HC2 | 4 |
| 9 | 2 | LC1-HC1 | 1 | 6 | LC2-HC2 | 4 |
| 10 | 3 | LC1-HC1 | 2 | 9 | LC2-HC2 | 4 |
| **co-transfections** |  |  |  |  |  |  |
| Construct a | 3 | LC1 | 1 | 7 | HC1 | 4 |
| Construct b | 3 | LC2 | 1 | 7 | HC2 | 4 |
| **Pool** | **Construct transfection ratios** |  |  |  |  |  |
| 11 | a:b=1:1 |  |  |  |  |  |
| 14 | a:b=1:2 |  |  |  |  |  |
| 15 | a:b=2:1 |  |  |  |  |  |

### Cell culture

The cell culture media and the cell culture unit operations were performed as described previously (Rajendran *et al*., 2021) except for the clone ranking step performed in 24 deep well plates and the stability study production runs conducted in 125 mL shake flasks.

The miCHO GS expression host is a CHO-K1 derivative line where the native hamster glutamine synthase gene was downregulated using ATUM’s proprietary Leap-In transposase-based technology. The stability of the GS knock-down geno/phenotype enables glutamine-free and antibiotic-free selection similar to commercially available GS knock-out lines.

Transfection was performed using the standard miCHO electroporation protocol, as described earlier (Rajendran *et al*, 2021). Briefly, the Emicizumab and Vanucizumab producing stable pools were established by transfecting miCHO GS cells with 25 μg total 2ORF, 3ORF, or 4ORF plasmid DNA with 3 μg Leap-In transposase mRNA. Following a 48-hour recovery period in non-selective media, the pools were selected under glutamine and antibiotic-free conditions.

### Protein purification

Emicizumab and Vanucizumab clarified harvests from the pool ranking and the stability production runs of the final clones were purified via affinity chromatography. The supernatant was loaded onto Protein A resin (Cytiva) that had been equilibrated with PBS, pH 7, and the resin was washed with PBS, pH 7. The proteins were eluted via isocratic elution with 50 mM Sodium Acetate, 100 mM NaCl, pH 3.4, and the eluates were neutralized with Tris base. The eluates were loaded onto Sephadex G-25 columns (Cytiva) for buffer exchange into PBS, pH 7.0.

### Analytical characterization

Productivity was assessed by biolayer interferometry. Clarified supernatants were diluted using Sample Diluent (Pall Forte Bio, cat#18-1048) and measured in 96-well plates on an Octet HTX (Pall Forte Bio; Sartorius) instrument using Protein A sensors (Sartorius, cat#18-5010).

Appropriate sample dilutions were generated, and all valid measurement values were within the linear standard concentration range. Various replicate measurements were also performed to reduce dilution and sensor-dependent measurement errors.

Droplet digital PCR (ddPCR) was utilized for stably integrated transgene copy number (CN) determination from the CHO cell substrates.

Bio-Rad’s QX200 Droplet Digital PCR System and ddPCR Supermix for Probes reagent (Catalog number 186-3027) were used for CN determinations in CHO cell genomic DNA prepared by the Zymo Research Quick DNA Miniprep Kit (Catalog number D3025). PrimeTime qPCR5’ nuclease probes and PCR primer sets were purchased from Integrated DNA Technology.

The raw data was analyzed by QuantaSoft™ Analysis Pro following the manufacturer’s standard protocol. The transgene-specific primer set was designed to amplify a unique sequence (not present in the CHO genome) from the glutamine synthetase element and used FAM as the reporter dye. The CHO-specific reference primer set was designed to amplify sequences from exon 3 of the Tmed2 gene and used HEX as reporter dye.

#### Heterodimer Quantitation Assay by cIEX HPLC

ProA purified material from Emicizumab expressing pools/clones was further analyzed by cIEX HPLC to assess the heterodimer fraction. ProA purified protein samples were diluted 1:1 with 50 mM Sodium Phosphate pH 6.0 and loaded on an Agilent Bio Mab NP5, SS 5 µm 4.6 × 250 mm column at a flow rate of 1ml/min. Protein fractions were eluted using a 6-8 pH gradient. Purified homodimer controls were used as standards to help assess the heterodimer fraction from pools and clone-derived material.

#### Heterodimer Quantitation Assay by HIC HPLC

ProA purified material from Vanucizumab expressing pools/clones was further analyzed by HIC HPLC to assess the heterodimer fraction. ProA purified protein samples were diluted 1:1 with 40 mM Sodium Phosphate, 3 M Ammonium Sulfate, pH 6.5, and loaded on a ProPac HIC-10 column (2.1 x 100 mm) at a flow rate of 0.2 mL/min. Protein fractions were eluted using a decreasing ammonium sulfate salt gradient. Purified homodimer controls were used as standards to help assess the heterodimer fraction of material derived from pools & clones.

Homodimer control samples (hole-hole and knob-knob) were expressed in stable pools and purified using proA resin. These controls were run alongside purified protein samples isolated from the stable pools and clones. To identify heterodimer peaks in the pools/clones, retention time matching was performed with peaks in homodimer controls. The matched peaks were labelled as hole-related contaminants or knob related contaminants. To quantify heterodimer peak %, the area of the peak of interest (heterodimer peak) was divided by the total area of all the identified peaks (heterodimer, hole-related contaminants, and knob related contaminants).

## Results

### Stable Pool Development

The viability curves during stable pool selection are shown in Figures 2a and 2b. All stable pools recovered within 14 days, the typical Leap-In transposase-mediated selection timeframe.

**Figure 2:**
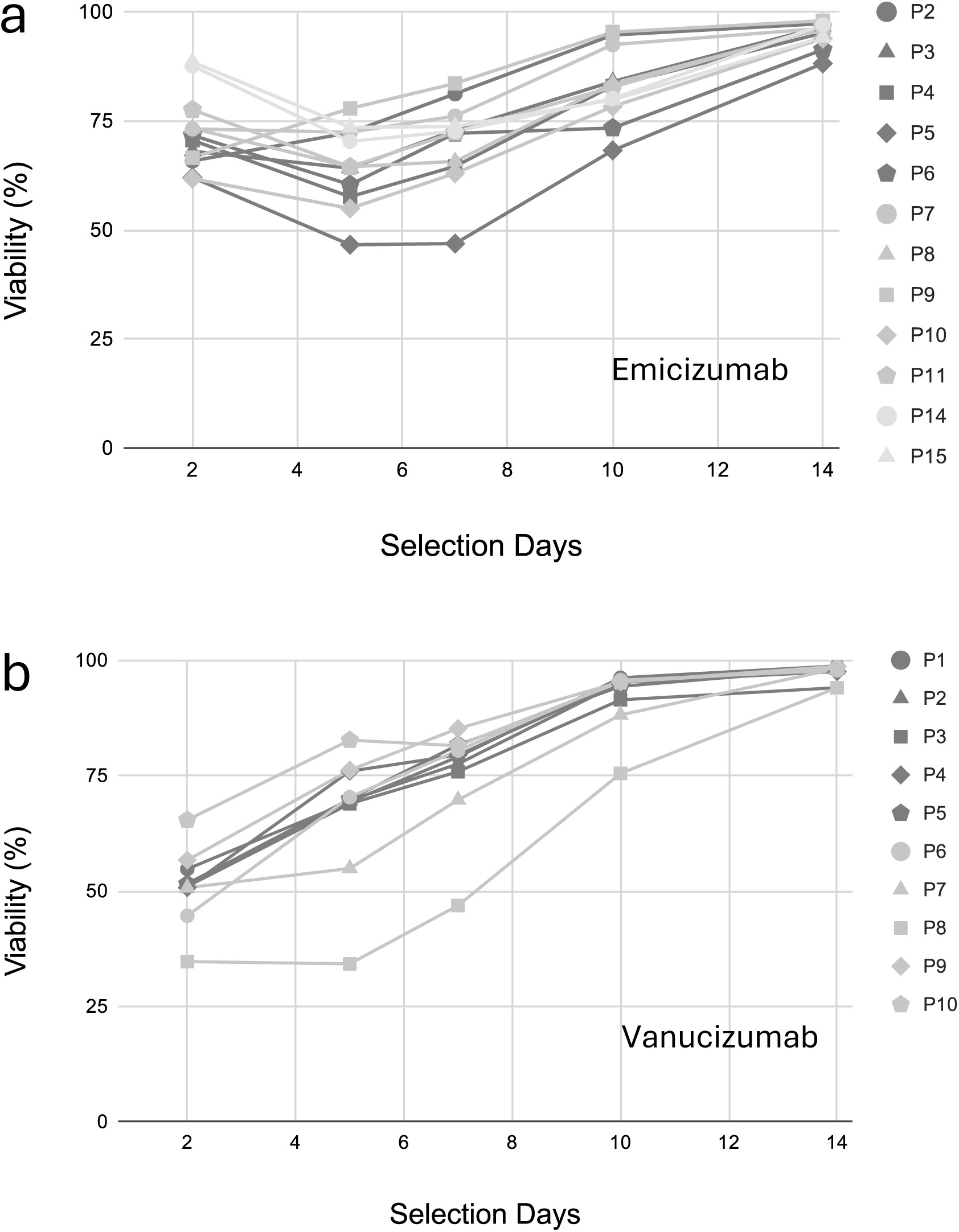
Recovery of the Emicizumab (a) and the Vanucizumab (b) pools during stable pool selection.

There was a difference between the viability profiles of the Emicizumab (Figure 2a) and the Vanucizumab (Figure 2b) pools during selection. The Emicizumab pools had lower viability (∼40-60%) at the end of the two-day non-selective recovery period compared to the Vanucizumab pools (60% to 80%). From that time point on, during the glutamine-free selection, the viability of the Emicizumab pools did not drop further, while most of the Vanucizumab pools decreased their viability. Also, out of the 3ORF Emicizumab pools, Pool 8 had the slowest recovery but highest productivity and high heterodimer fraction, contrasting with Vanucizumab Pool 5, where slow recovery was associated with low productivity and low heterodimer formation (Table 3a and 3b).

**Table 3.**
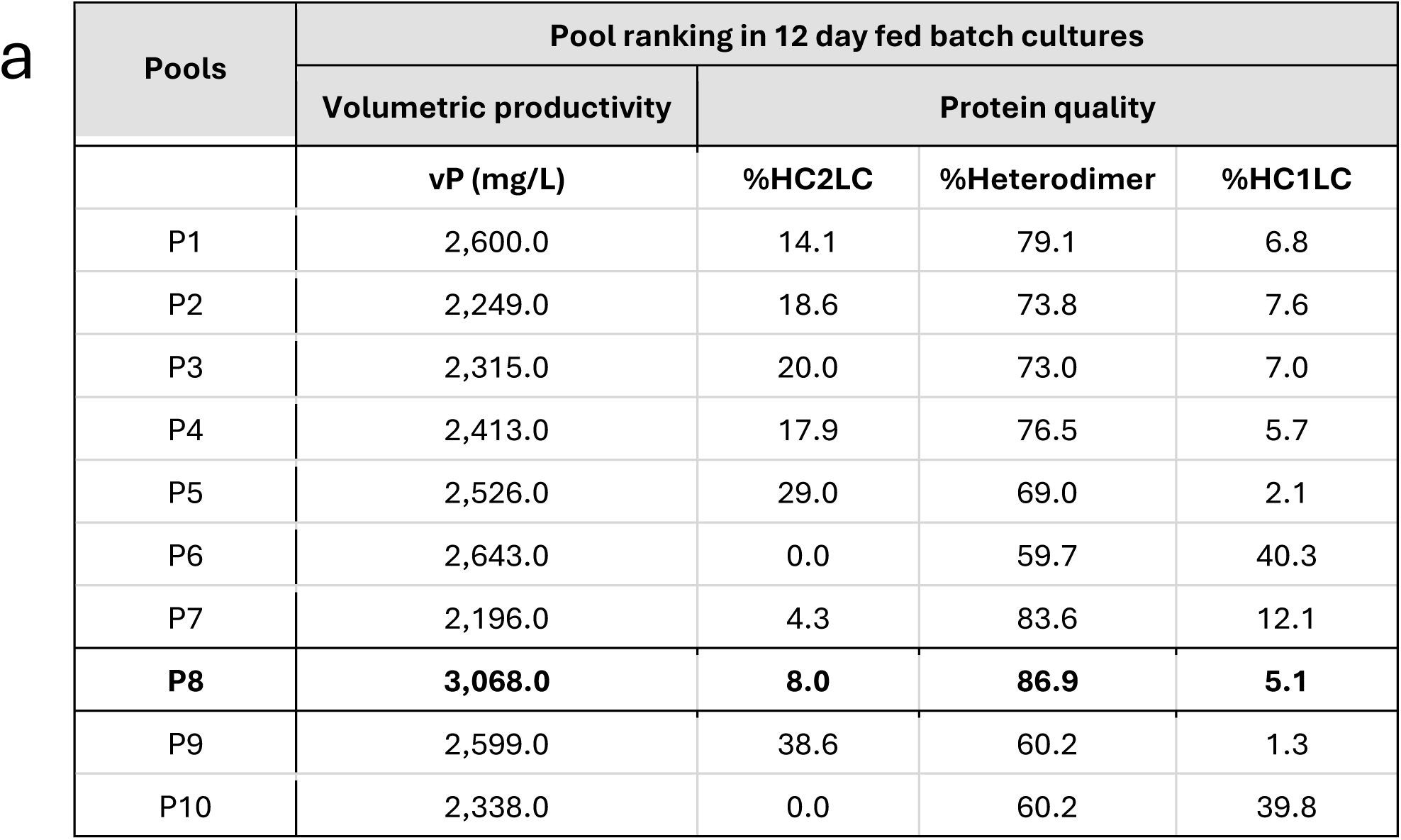
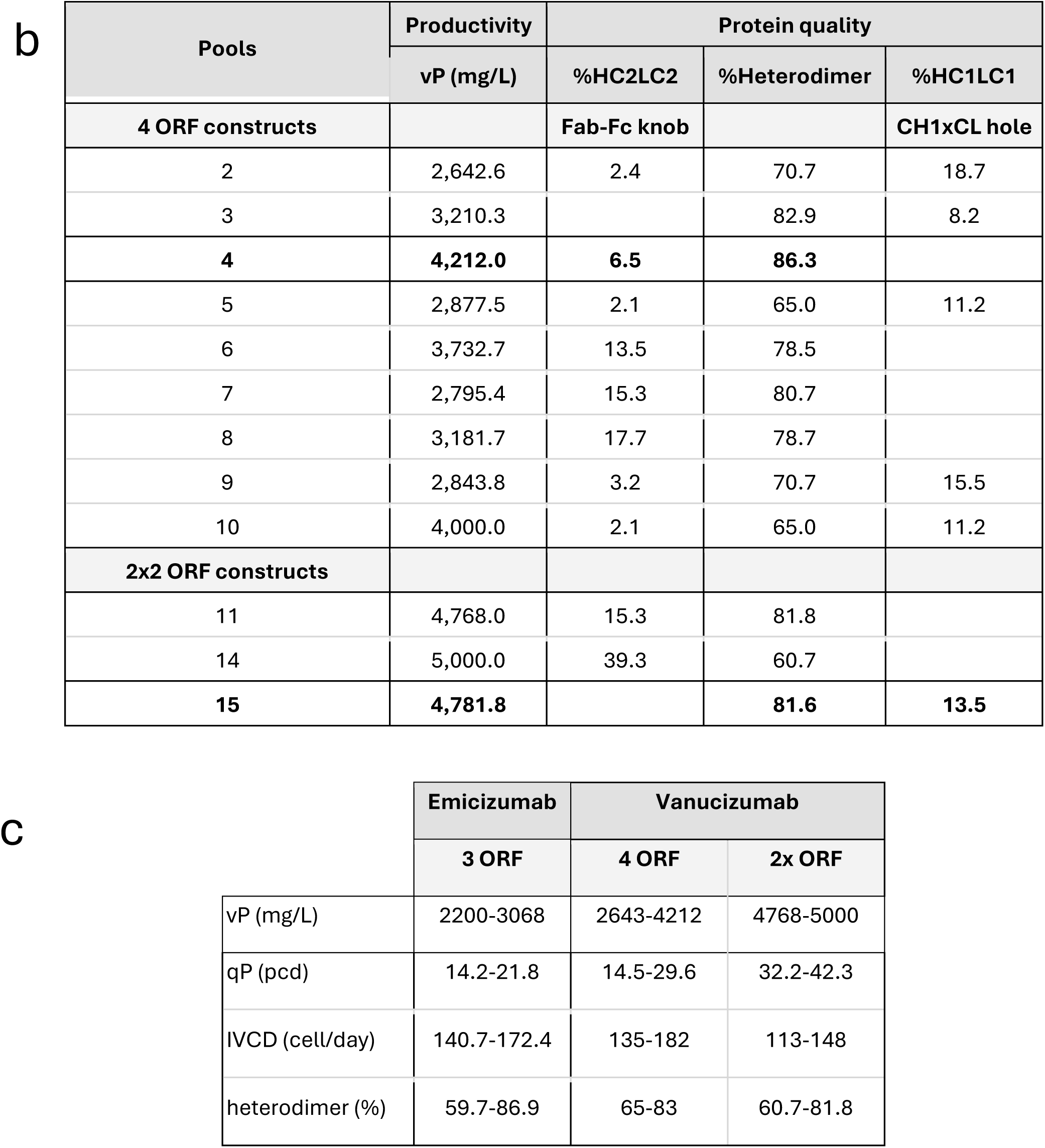
Volumetric productivity and heterodimer fraction in Emicizumab (a) and Vanucizumab (b) pools. Performance parameter ranges of the Emicizumab and Vanucizumab pools (c).

| Pools | Pool ranking in 12 day fed batch cultures |  |  |  |
| --- | --- | --- | --- | --- |
|  | Volumetric productivity | Protein quality |  |  |
|  | vP (mg/L) | %HC2LC | %Heterodimer | %HC1LC |
| P1 | 2,600.0 | 14.1 | 79.1 | 6.8 |
| P2 | 2,249.0 | 18.6 | 73.8 | 7.6 |
| P3 | 2,315.0 | 20.0 | 73.0 | 7.0 |
| P4 | 2,413.0 | 17.9 | 76.5 | 5.7 |
| P5 | 2,526.0 | 29.0 | 69.0 | 2.1 |
| P6 | 2,643.0 | 0.0 | 59.7 | 40.3 |
| P7 | 2,196.0 | 4.3 | 83.6 | 12.1 |
| **P8** | **3,068.0** | **8.0** | **86.9** | **5.1** |
| P9 | 2,599.0 | 38.6 | 60.2 | 1.3 |
| P10 | 2,338.0 | 0.0 | 60.2 | 39.8 |

b
| Pools | Productivity | Protein quality |  |  |
| --- | --- | --- | --- | --- |
|  | vP (mg/L) | %HC2LC2 | %Heterodimer | %HC1LC1 |
| **4 ORF constructs** |  | **Fab-Fc knob** |  | **CH1xCL hole** |
| 2 | 2,642.6 | 2.4 | 70.7 | 18.7 |
| 3 | 3,210.3 |  | 82.9 | 8.2 |
| **4** | **4,212.0** | **6.5** | **86.3** |  |
| 5 | 2,877.5 | 2.1 | 65.0 | 11.2 |
| 6 | 3,732.7 | 13.5 | 78.5 |  |
| 7 | 2,795.4 | 15.3 | 80.7 |  |
| 8 | 3,181.7 | 17.7 | 78.7 |  |
| 9 | 2,843.8 | 3.2 | 70.7 | 15.5 |
| 10 | 4,000.0 | 2.1 | 65.0 | 11.2 |
| **2x2 ORF constructs** |  |  |  |  |
| 11 | 4,768.0 | 15.3 | 81.8 |  |
| 14 | 5,000.0 | 39.3 | 60.7 |  |
| **15** | **4,781.8** |  | **81.6** | **13.5** |

**Table 3**
|  | Emicizumab | Vanucizumab |  |
| --- | --- | --- | --- |
|  | 3 ORF | 4 ORF | 2x ORF |
| vP (mg/L) | 2200-3068 | 2643-4212 | 4768-5000 |
| qP (pcd) | 14.2-21.8 | 14.5-29.6 | 32.2-42.3 |
| IVCD (cell/day) | 140.7-172.4 | 135-182 | 113-148 |
| heterodimer (%) | 59.7-86.9 | 65-83 | 60.7-81.8 |

### Stable Pool Ranking

The Emicizumab and the Vanucizumab pools were ranked using 12 or 14-day fed batch cultures, respectively. The ranking parameters were volumetric and specific productivities of the cultures and heterodimer fractions in the purified protein batches. The viable cell density (VCD) and the viability profiles of the fed batch cultures are shown in Figures 3a-d.

**Figure 3:**
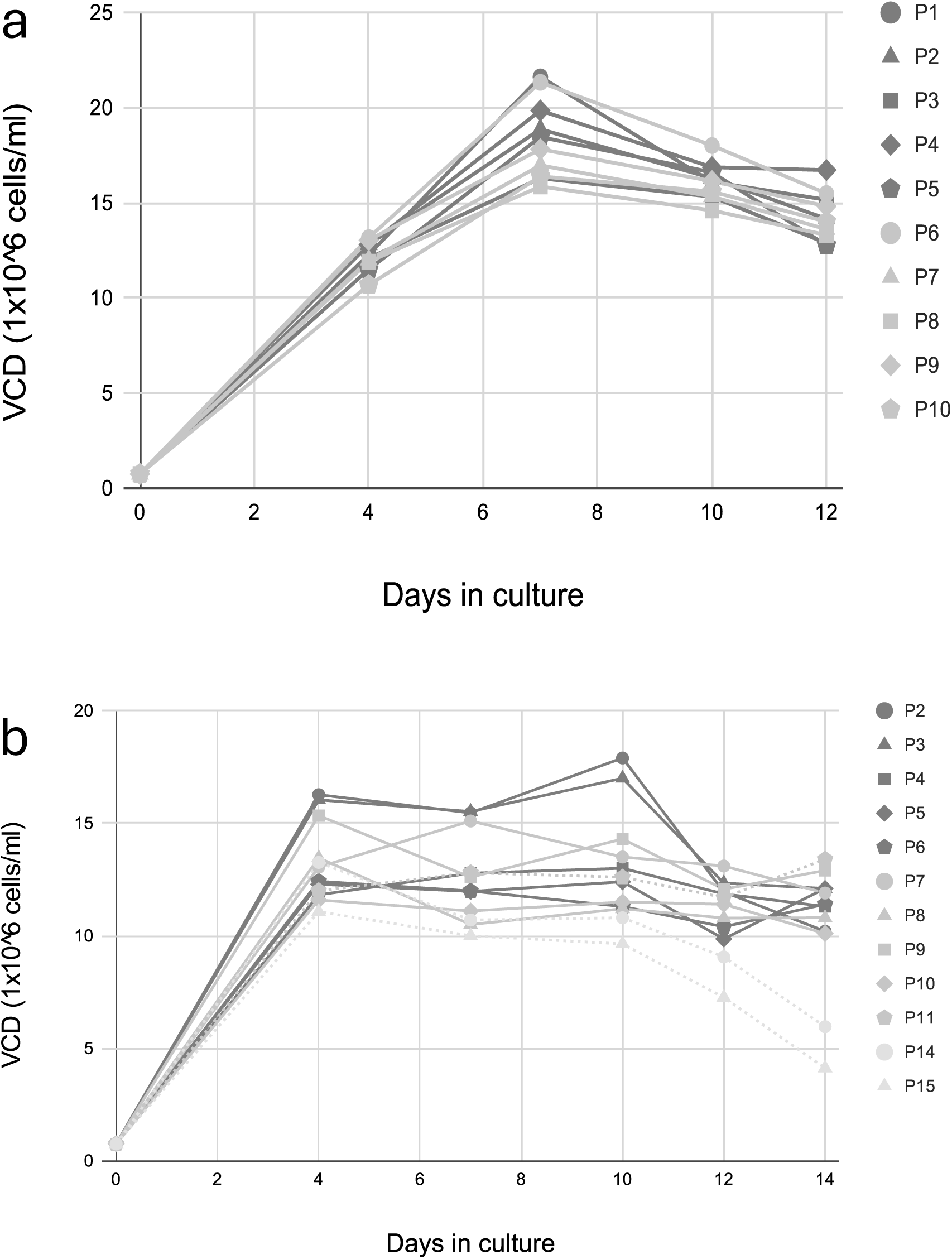
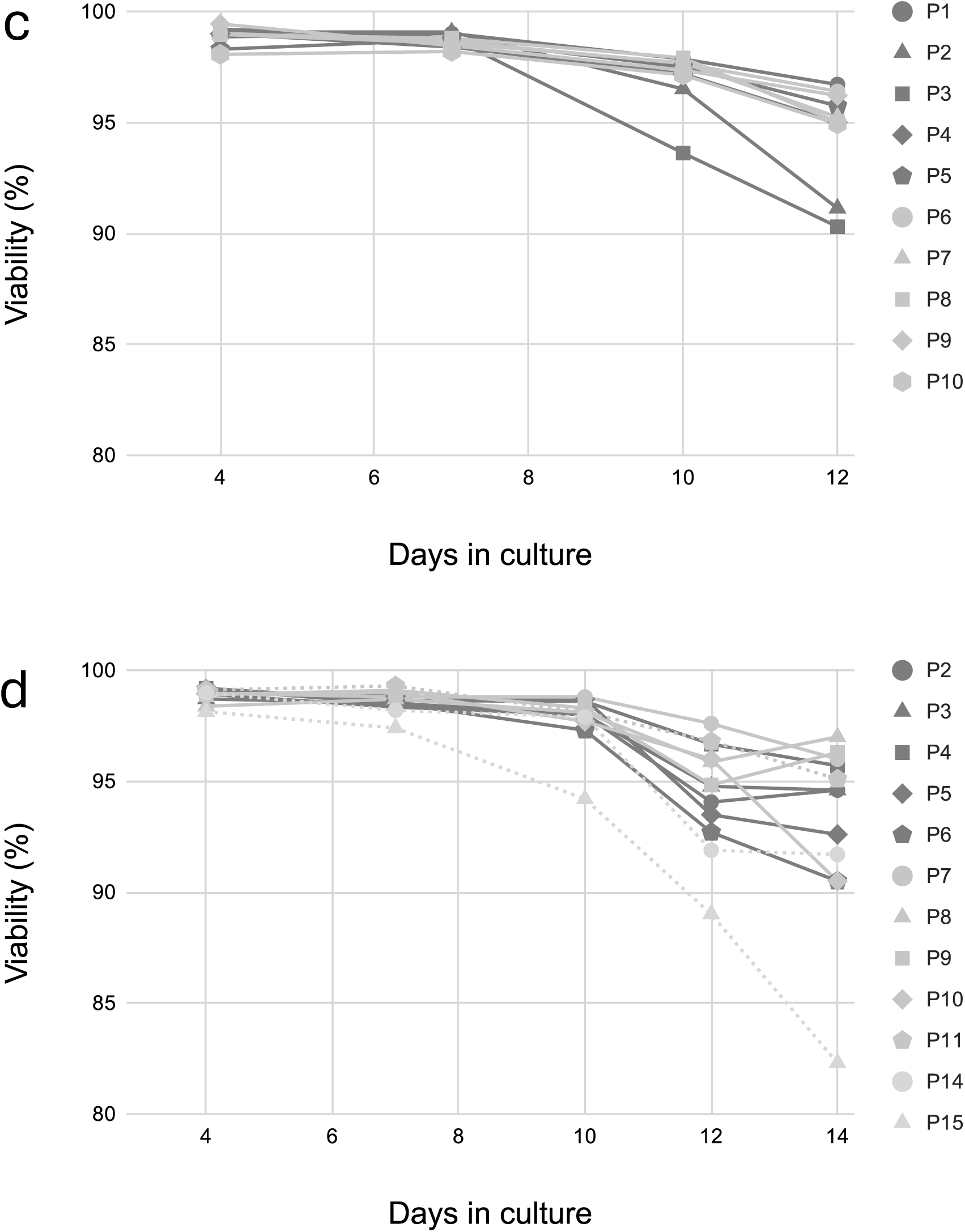
VCD and viability profiles of the Emicizumab and the Vanucizumab stable pools during the pool ranking production runs. Emicizumab pools VCD (a), Vanucizumab pools VCD (b), Emicizumab pools viability (c), Vanucizumab pools viability (d).

The majority of the Emicizumab pools reached higher peak VCDs compared to the Vanucizumab pool cultures. Day 12 viabilities in the 3ORF Emicizumab pools were moderately higher compared to the four chain pools on day 12. The pools transfected with single 3ORF or single 4ORF constructs maintained higher than 90% viability on day 12 or day 14, respectively. In the Vanucizumab co-transfection pools, the end viabilities were influenced by the different plasmid ratios (LC2 and HC2): (LC1 and HC1), with the 1:1 ratio at 96%, the 2:1 ratio at 92% and the 1:2 ratio at 82%.

The productivity and heterodimer fraction data are presented in Tables 3a and b, and a comparison of the ranking parameter ranges is shown in Table 3c.

Considering the different molecular designs and subunit sequence differences between Emicizumab and Vanucizumab, the direct comparison of the 3ORF and 4ORF pools is not straightforward.

On the other hand, there were significant differences between the volumetric and specific productivities as well as the IVCDs of the 4ORF and the 2 x 2ORF Vanucizumab pools. As expected, the qP and the IVCD ranges were inversely correlated. The productivity differences did not affect heterodimer fractions, and in general, no significant correlation could be found between the purified heterodimer fraction and the vP or the qP of the pools (not shown). Pool 8 from the Emicizumab set, Pool 4 from the 4ORF transfection, and Pool 15 from the co-transfection Vanucizumab sets were selected for single cell cloning.

### Clone Ranking and Clonal Productivity Distribution

The clones were ranked by productivity and growth in seven-day fed batch cultures in 24 deep well plates. The clonal volumetric productivity distributions are presented in Figures 4a, b, and c.

**Figure 4:**
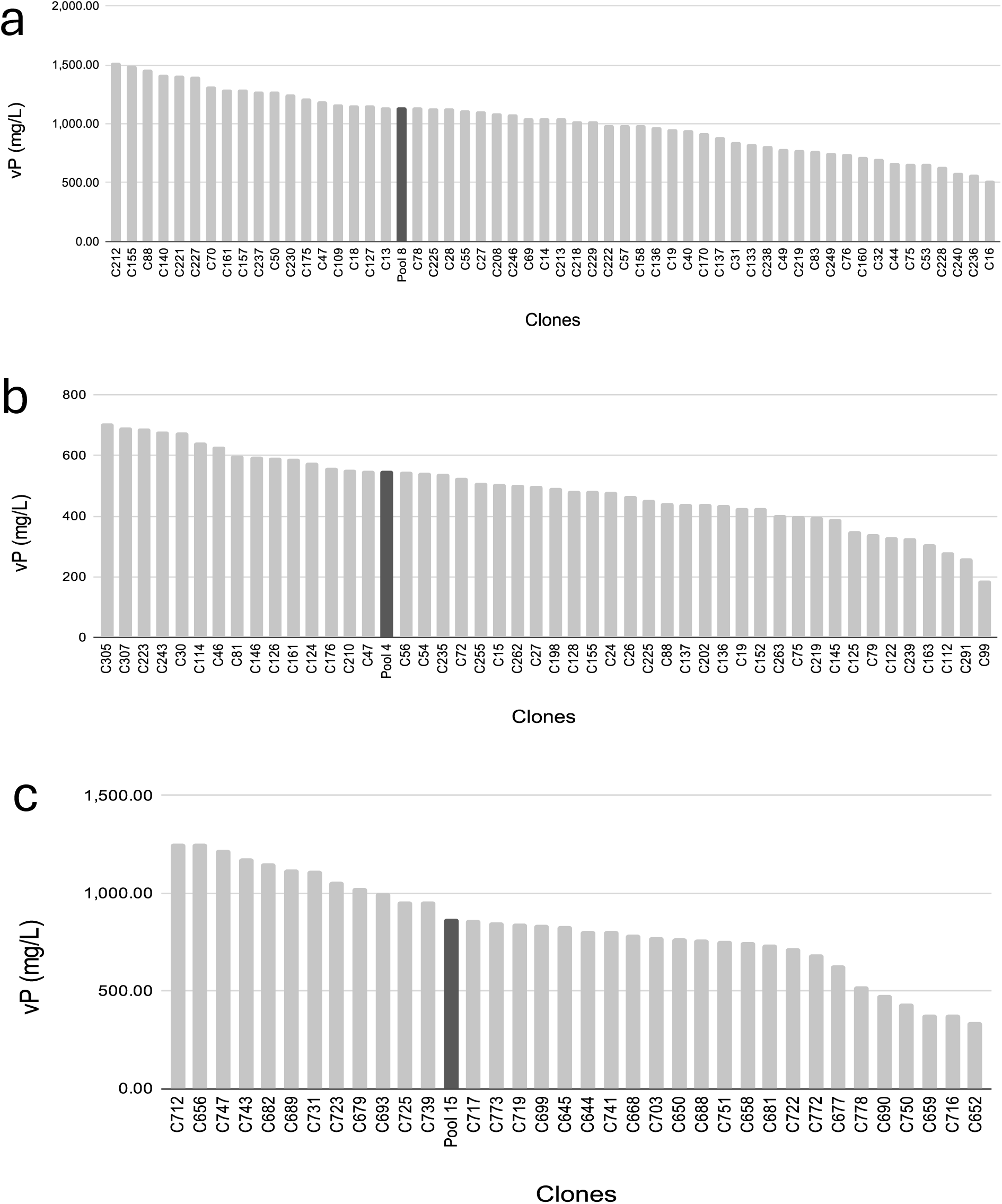
Parent pools’ volumetric productivities and the corresponding clonal volumetric productivity distributions. 3ORF Emicizumab Pool 8 (a), 4ORF Vancizumab Pool 4 (b), 2×2ORF co-transfected Vanucizumab Pool 15 (c).

The clonal distributions represent the typical Leap-In mediated profile with >30% of the clones in the first and >80% of the clones in the first and second productivity quartiles. There were no clones in the fourth quartile, and indeed, no non-producers were observed (Table 4, lower panel).

**Table 4.**
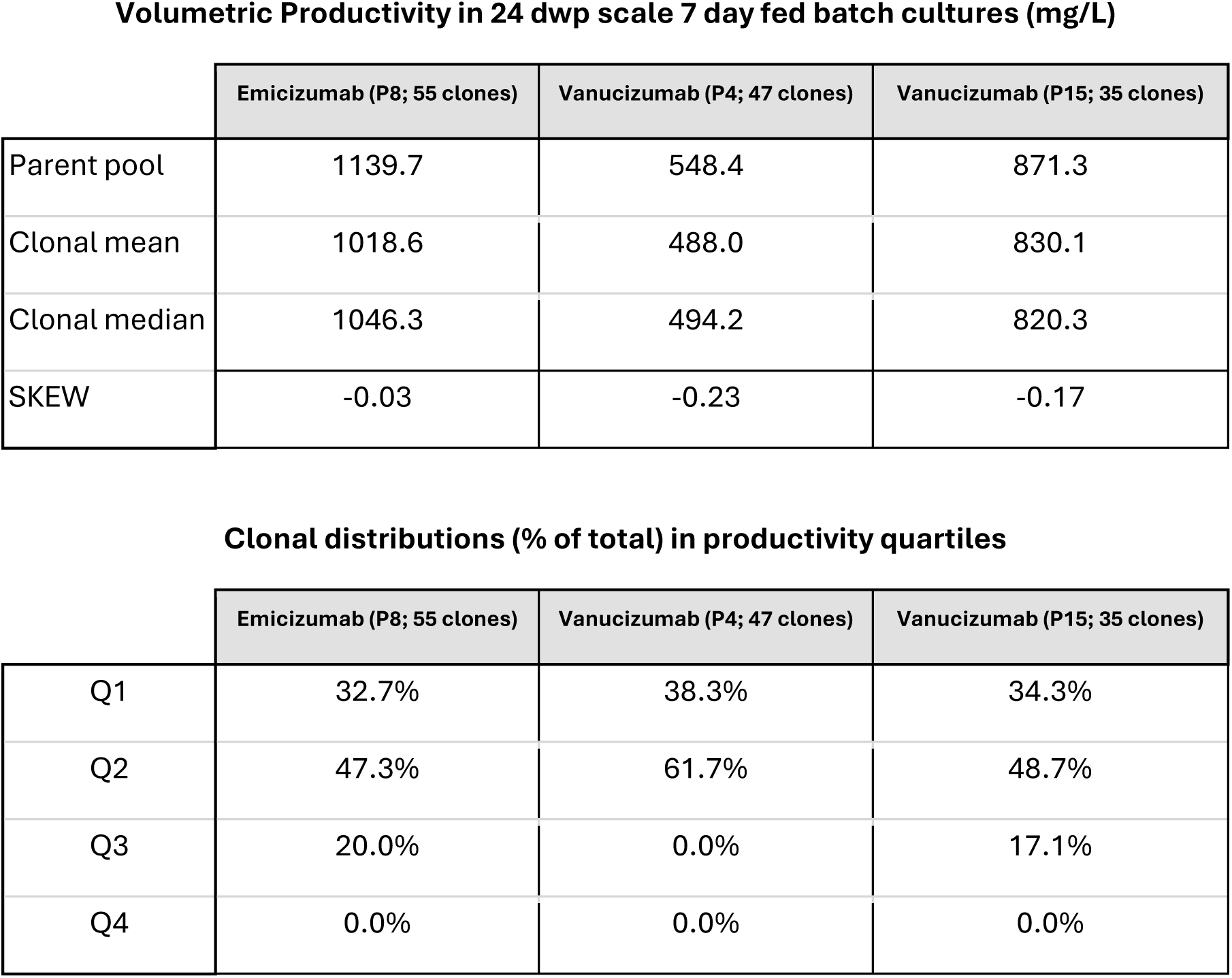
Upper panel: Statistical characterization of clonal distributions and pool-to-clone volumetric productivity comparability in the three selected pools (Pool 8, Pool 4, and Pool 15) and their corresponding derivative clones. Lower panel: Sorting the stable clones into volumetric productivity quartiles.

Volumetric Productivity in 24 dwp scale 7 day fed batch cultures (mg/L)
|  | Emicizumab (P8; 55 clones) | Vanucizumab (P4; 47 clones) | Vanucizumab (P15; 35 clones) |
| --- | --- | --- | --- |
| Parent pool | 1139.7 | 548.4 | 871.3 |
| Clonal mean | 1018.6 | 488.0 | 830.1 |
| Clonal median | 1046.3 | 494.2 | 820.3 |
| SKEW | -0.03 | -0.23 | -0.17 |

**Table 4**
|  | Emicizumab (P8; 55 clones) | Vanucizumab (P4; 47 clones) | Vanucizumab (P15; 35 clones) |
| --- | --- | --- | --- |
| Q1 | 32.7% | 38.3% | 34.3% |
| Q2 | 47.3% | 61.7% | 48.7% |
| Q3 | 20.0% | 0.0% | 17.1% |
| Q4 | 0.0% | 0.0% | 0.0% |

The comparison of pool productivity to the derivative clonal mean and median productivity values demonstrates that the three parameters are very close in all three data sets. This observation, together with the calculated SKEW values, indicates that the clonal productivity distributions in Leap-In mediated stable pools expressing bispecific antibodies are balanced without any significant skew in any direction (Table 4, upper panel). These characteristics are in agreement with the practical observations that a relatively small number—in this study 35, 47, and 55—of clones is sufficient to identify high producer stable clones expressing protein with desired quality attributes, in this case, high heterodimer fraction.

Based on their productivity and growth characteristics, a subset of clones was selected from each pool for stability studies and heterodimer fraction assessment in 14-day 25 ml scale shake flask cultures. Nine clones from Emicizumab Pool 8, eight clones from Vanucizumab Pool 4, and nine clones from Vanucizumab Pool 15 moved forward.

### Comparability and Stability Assessment

The Emicizumab-producing 3ORF Pool 8, the Vanucizumab-producing 4ORF Pool 4, and the Vanucizumab-producing 2 x 2ORF Pool 15, together with their respective selected clones, have been passaged up to 60 population doublings (PD) in glutamine-free medium. The stable clones were also passaged in media supplemented with 5 mM glutamine. The Time 0 (T0) and the PD60 cultures from the three pools and the derivative clones were cryopreserved. 14-day fed batch cultures were set up from the archived cultures. The most critical product quality attributes, volumetric productivity and heterodimer fraction, were measured in the Day 14 harvests. To assess stability and pool-to-clone comparability, average clonal volumetric productivities and heterodimer fractions were calculated and graphed for the T0 and PD60 timepoints. In addition to the actual volumetric productivities and heterodimer fractions, the measurements at PD60 are also presented as % of the corresponding T0 values.

### Emicizumab expressed by a 3ORF transposon

#### Volumetric Productivity

The volumetric productivity data, from Pool 8 and its derivative clones, are presented in Figures 5a-c.

**Figure 5:**
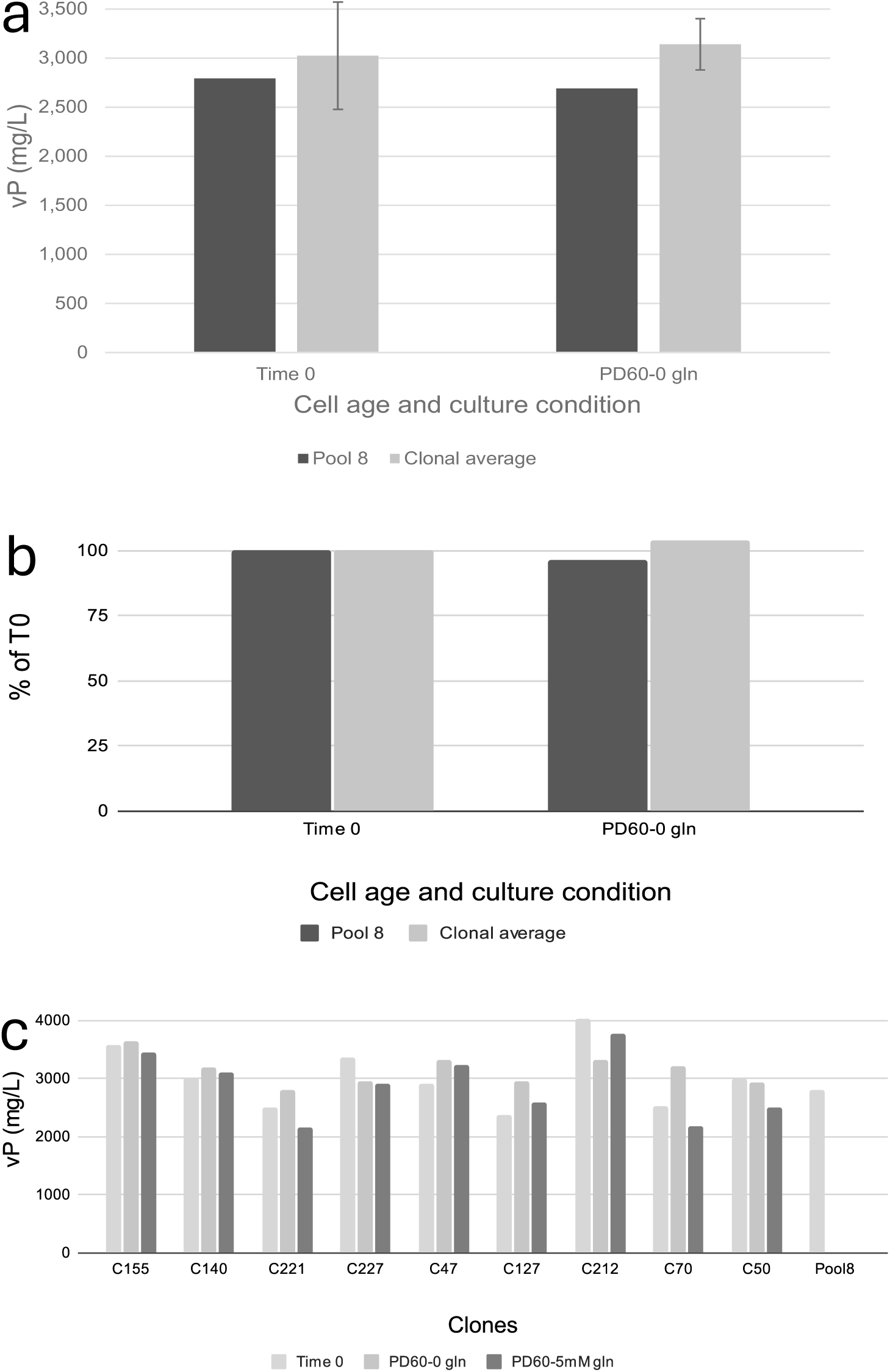
Volumetric productivity comparability and stability in the Emicizumab expressing Pool 8 and its derivative clones. Pool 8 and clonal average volumetric productivities at T0 and PD60 (a). PD60 Pool 8 and clonal average volumetric productivities expressed as % of the corresponding T0 values (b). Clonal volumetric productivity, stability, and comparison of clonal productivity to the T0 Pool 8 value (c).

The results demonstrate that the PD60 clonal averages were maintained at the T0 level, and even Pool 8 PD60 volumetric productivity remained at 96% of the T0 level when passaged under selective pressure. (Figure 5a, b.) This data indicates that the volumetric productivity of the individual clones derived from Pool 8 is stable. To further confirm clonal productivity stability, all clones met the industry rule of thumb (Hertel *et al*. 2022), “PD60 productivity >70% of T0 productivity” stability cut off (Figure 5c).

The data also demonstrates that the productivity of Pool 8 at T0 is highly comparable to the average productivity of its derivative clones and extends the Leap-In mediated stable pools’ productivity predictability to heterodimeric bispecific molecules (Figures 5a, b). This predictability is further exemplified by comparing the individual derivative clonal productivities with the source Pool 8 productivities (Figure 5c)

#### Heterodimer fraction

Day 14 harvests were clarified and purified, and the heterodimer fractions were measured as described in the Materials and Methods. The data is presented in Figures 6a-c.

**Figure 6:**
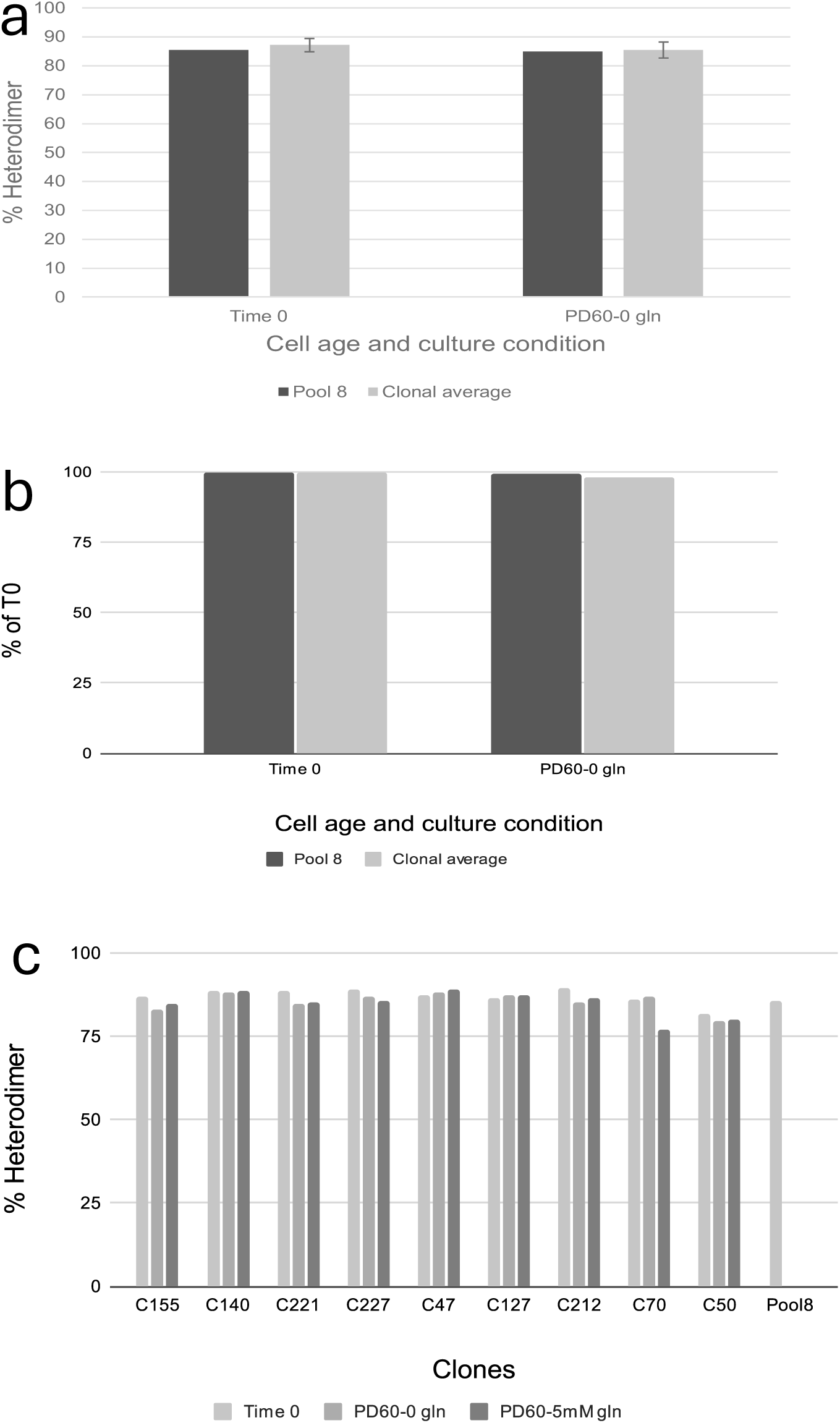
Heterodimer fraction comparability and stability in the Emicizumab expressing Pool 8 and its derivative clones. Pool 8 and clonal average heterodimer fractions at T0 and PD60 (a). PD60 Pool 8 and clonal average heterodimer fractions expressed as % of the corresponding T0 values (b). Clonal heterodimer fraction stability and comparison of the clonal heterodimer fractions to the T0 Pool 8 value (c).

The data in Figures 6a-c demonstrate that the pool to clone comparability and stability is consistent with the heterodimer fractions produced by the Leap-In mediated stable pools and clones. Figure 6a presents the heterodimer content at T0 and PD60 samples in both the pool-derived protein and in the derivative clones using the clonal average value. The absolute values are similar, demonstrating two advantages enabled by the Leap-In transposase system. The first is that the heterodimer fractions are highly comparable between the T0 Pool 8 and the T0 clonal average, indicating that the Leap-In stable pools predict the heterodimer fraction in their derivative clones (Fig.6a, 6c).

The second feature is the stability of the heterodimer fraction. The heterodimer content remains stable even in Pool 8 samples, where productivity decreased with pool age. The % decrease in heterodimer content is negligible in all samples (Fig. 5b and Fig. 6b). The heterodimer content remained stable in the individual clonal samples even when passaged in the presence of 5 mM glutamine (Figure 6c).

C70 PD60-5 mM glutamine sample was the only clonal example for non-perfect heterodimer stability. The heterodimer fraction decreased from 87% to 77%.

### Vanucizumab expressed by a 4ORF and a 2 x 2ORF Leap-In transposon design

#### Volumetric productivity

The Vanucizumab 4ORF and 2 x 2ORF pools and their derivative clones’ productivity data are presented in Figures 7a-f. The stable pools decreased their productivity along with aging, while the clonal productivities remained stable. The 4ORF Pool 4, passaged in the absence of glutamine, decreased productivity to 50% of the T0 value (Figures 7a and 7c). In contrast, Pool 15, established by co-transfection, maintained its productivity better than the 4ORF Pool 4, 64% of T0 (Figures 7 b and 7d).

**Figure 7:**
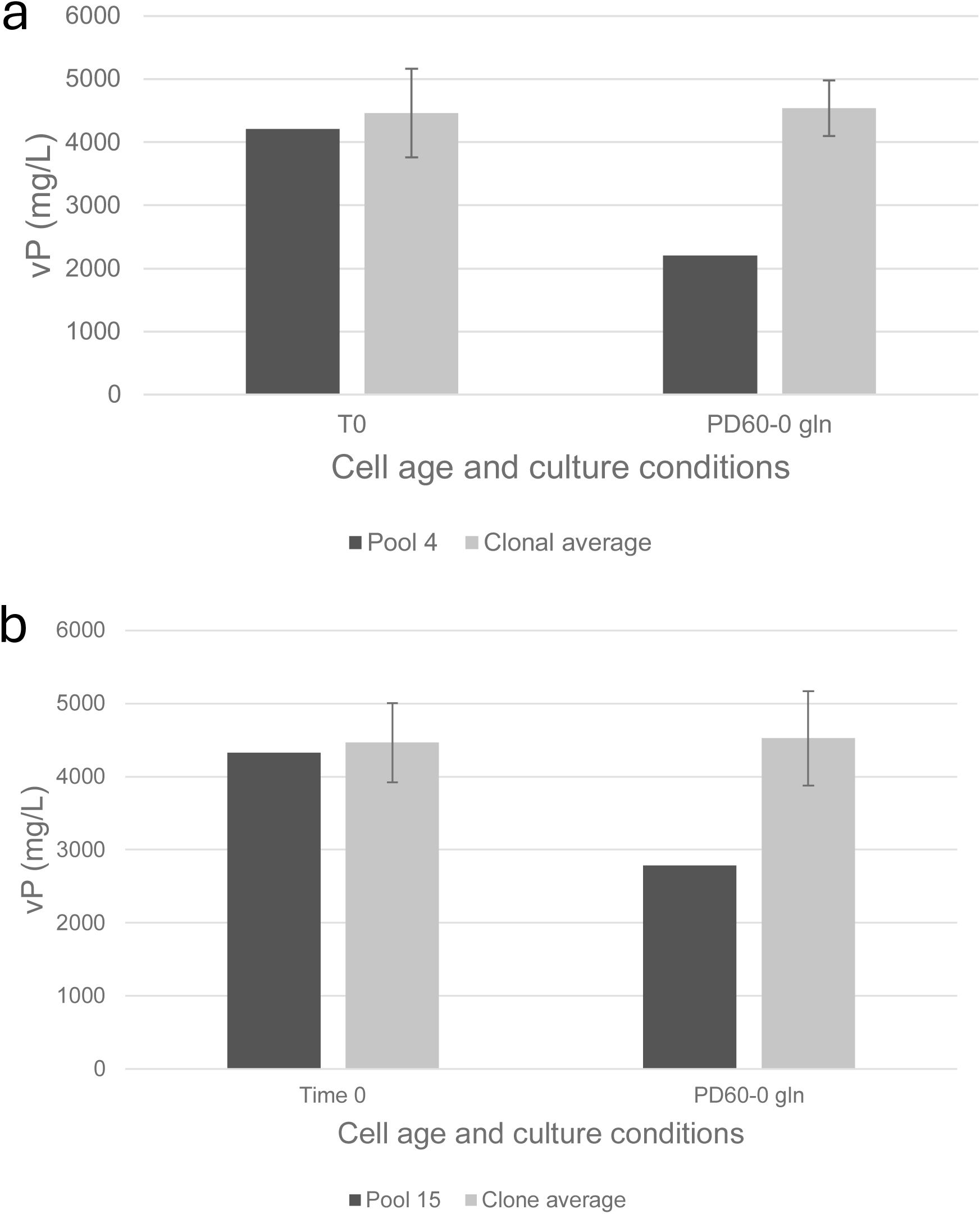
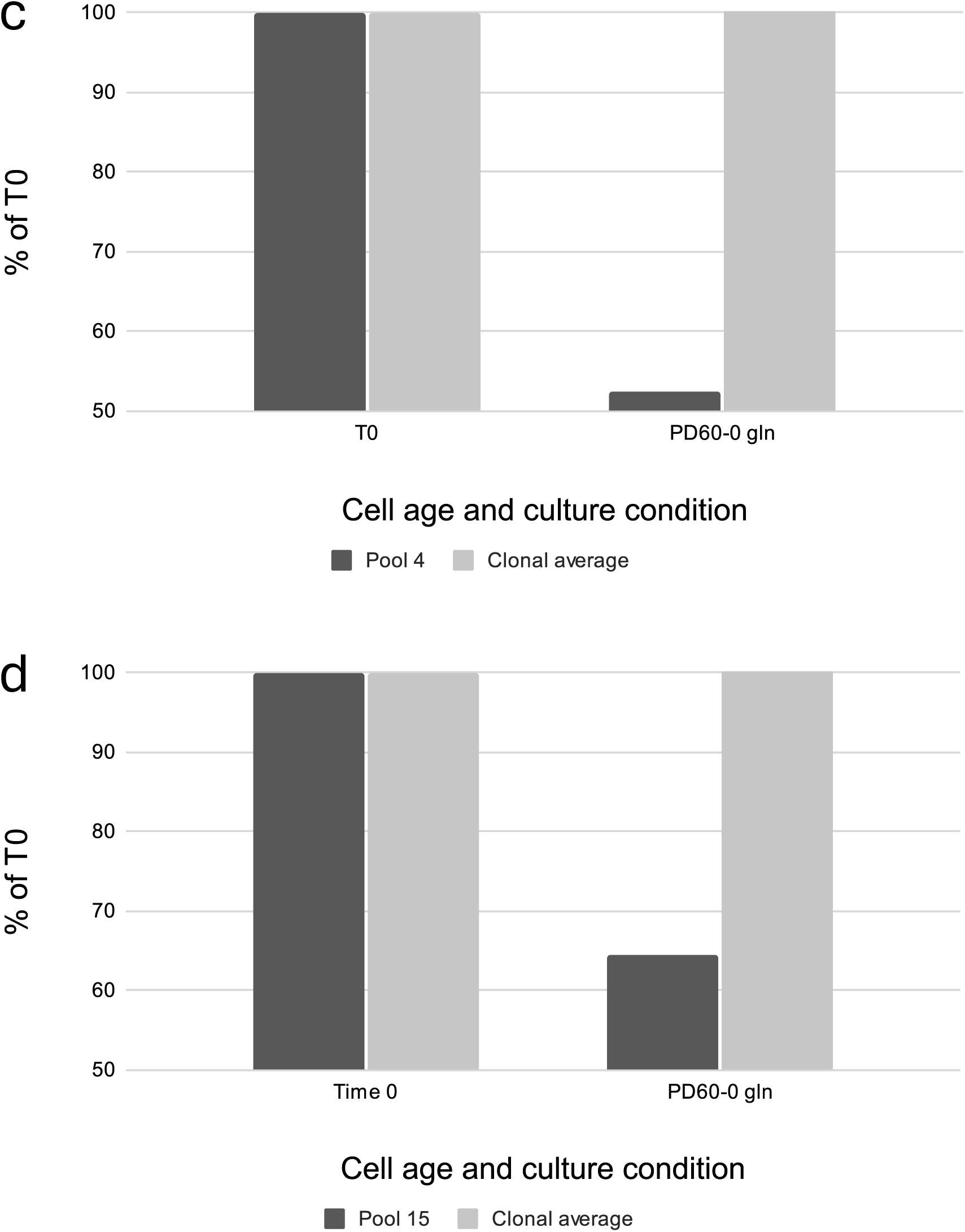
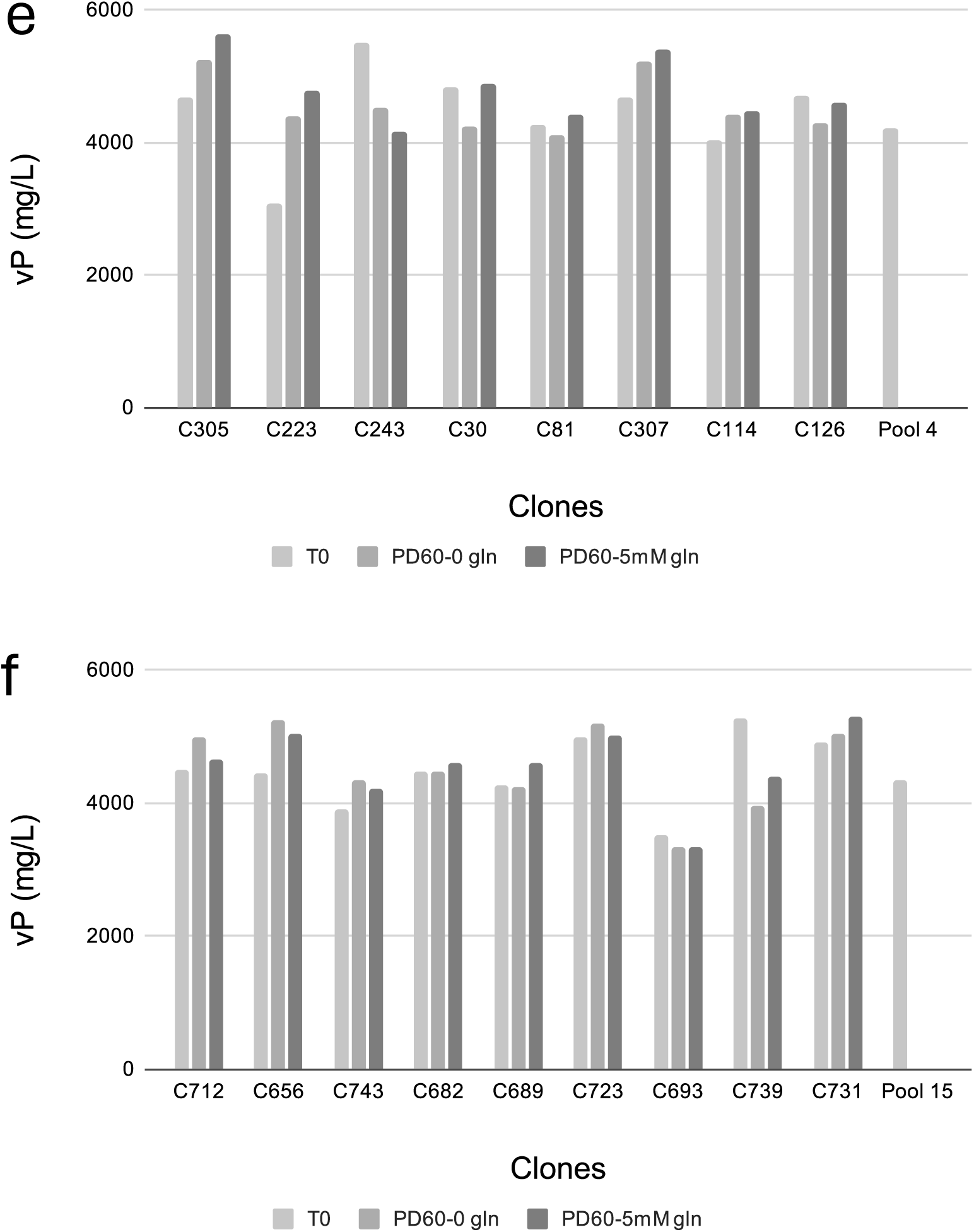
Volumetric productivity comparability and stability in the Vanucizumab expressing stable Pools 4 and 15 and their derivative clones. Pool 4 (a) and Pool 15 (b) and their respective clones’ average volumetric productivity at T0 and PD60. PD60 Pool 4 (c) and Pool 15 (d) and their respective clones’ average volumetric productivities expressed as % of the corresponding T0 values. Clonal volumetric productivity, stability, and comparison of the clonal productivities to the corresponding T0 Pool 4 (e) and Pool 15 (f) values.

The clonal productivity averages remained stable (Figures 7a and 7b). This indirect indication of clonal productivity stability is confirmed by data shown in Figures 7e and 7f, where all clones meet the industry standard productivity stability criteria.

The 4ORF and the 2 x 2ORF designs resulted in pools with similar volumetric productivities. The T0 pool and the derivative clonal average productivities were highly comparable in both Pool 4 and Pool 15, indicating that the final stable clone productivities are reliably predicted by the Leap-In-mediated parental pool productivity for this 4-chain bispecific antibody (Figures 7a and 7b).

#### Heterodimer fraction

The Vanucizumab 4ORF and 2 x 2ORF pools and their derivative clones’ heterodimer fraction data are presented in Figures 8a-f. In contrast to productivity, the heterodimer fractions remained stable in both pools at PD60. The clonal heterodimer fractions also remained stable, as shown by the individual clones in Figures 8e and f. The heterodimer fractions, detected in the two pools and in the corresponding clones represented by the clonal averages, were comparable (Figures 8a and 8b), thus supporting the concept of using the Leap-In pool to predict clonal heterodimer fraction comparability for 4-chain bispecific antibodies.

**Figure 8:**
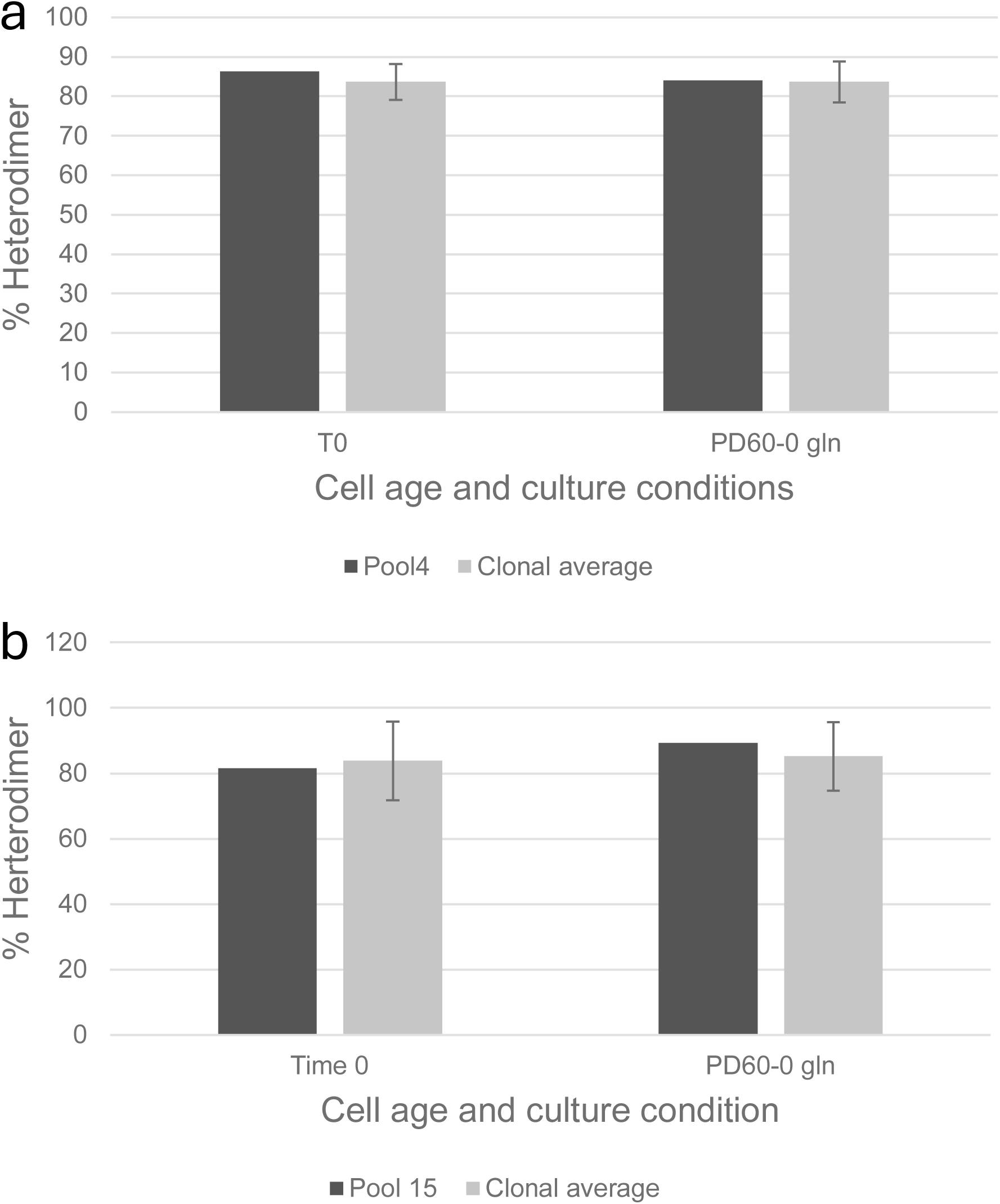
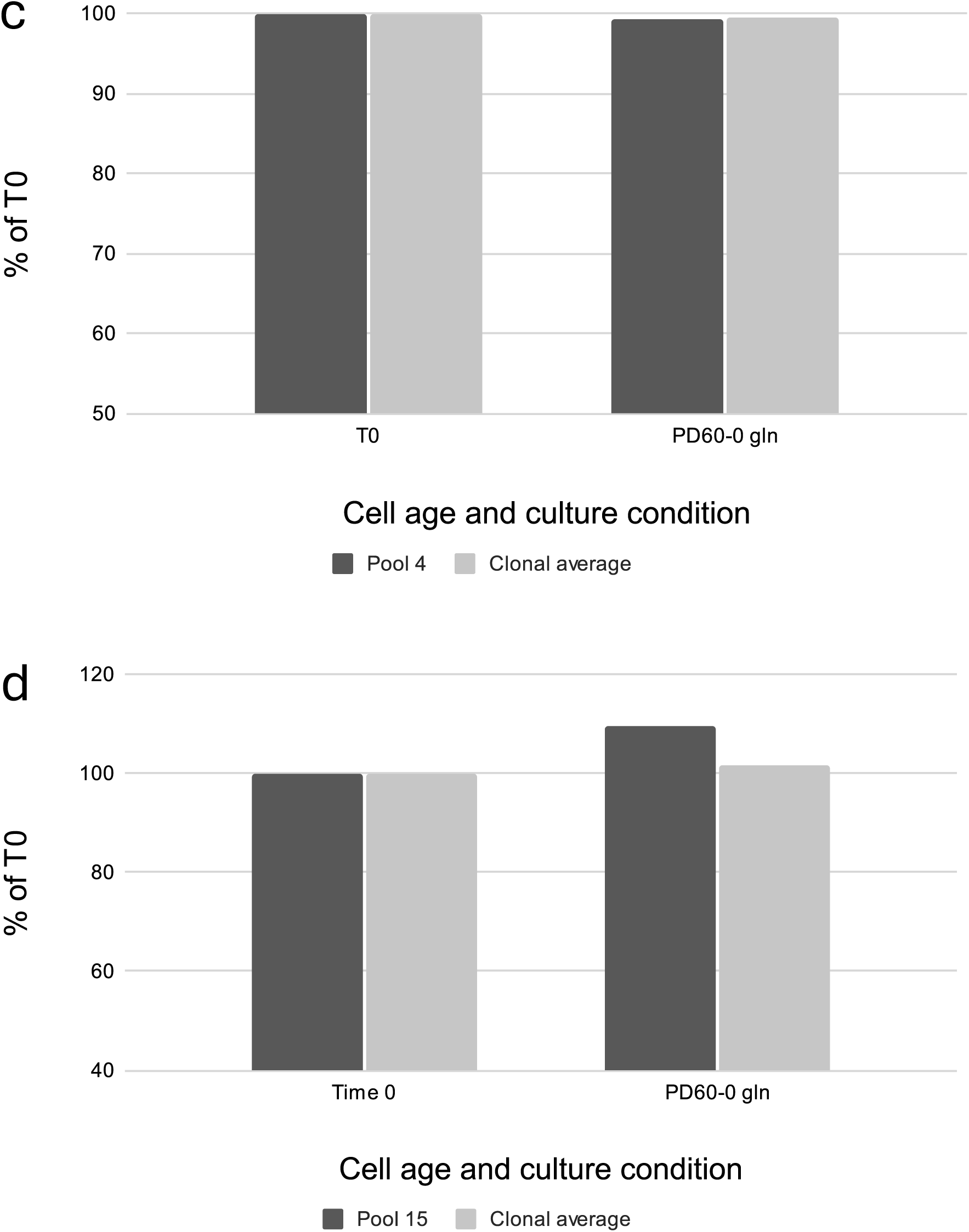
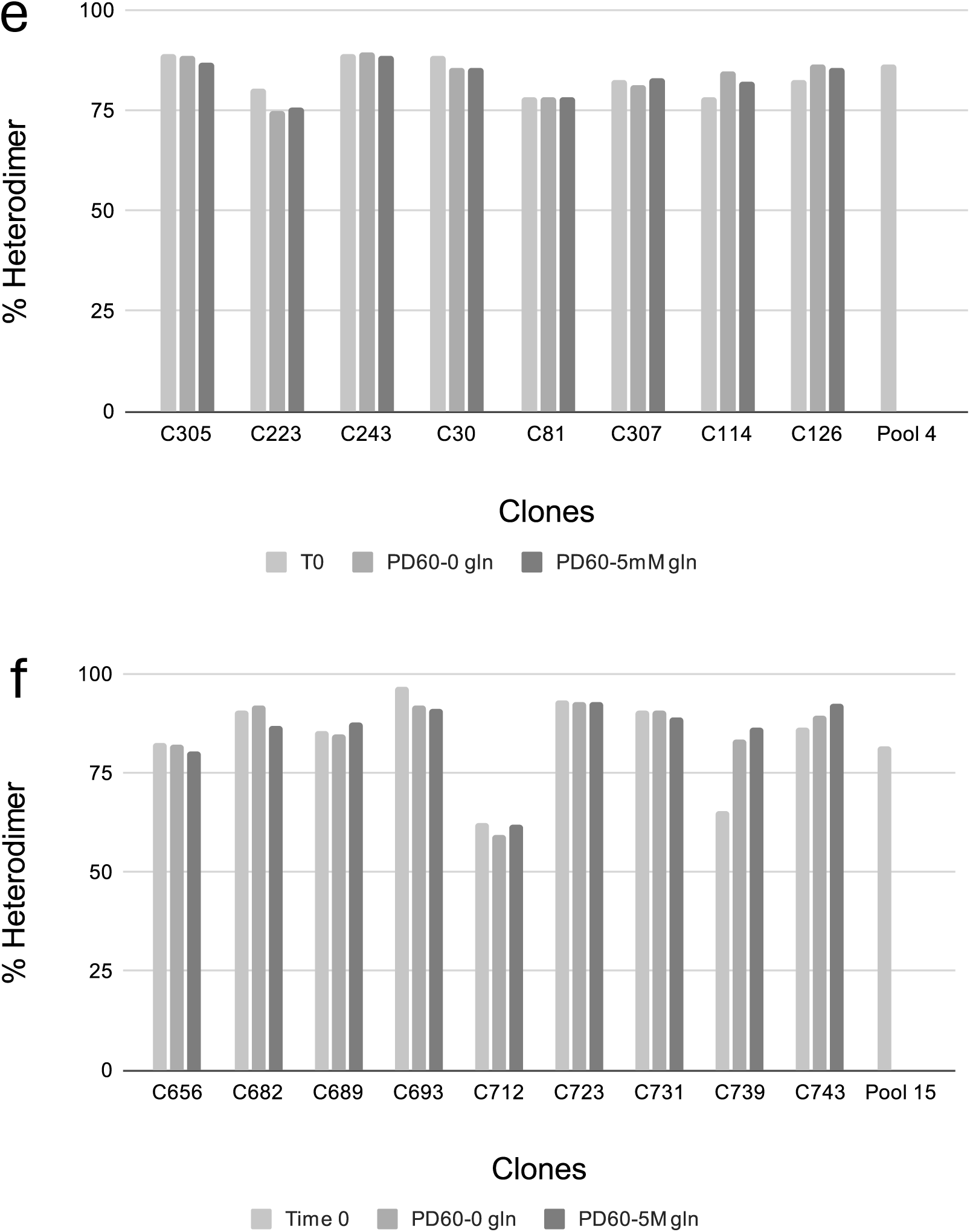
Heterodimer fraction comparability and stability in the Vanucizumab expressing stable Pools 4 and 15 and in their derivative clones. Pool 4 (a) and Pool 15 (b) and their respective clones’ average heterodimer fractions at T0 and PD60. PD60 Pool 4 (c) and Pool 15 (d) and their respective clones’ average heterodimer fractions expressed as % of the corresponding T0 values. Clonal heterodimer fraction stability and comparison of the clonal heterodimer fractions to the corresponding T0 Pool 4 (e) and Pool 15 (f) values.

#### Genomic copy number determination

The number of stably integrated Leap-In transposon-based expression constructs was determined using recombinant glutamine synthetase (GS) specific reagents. The target sequence is present at one copy/transposon in all three (3ORF, 4ORF, and 2ORF) types of constructs. The measurements were performed in the stable pools and the derivative clones at T0 and at the end of the PD60 stability passages.

The data is presented in Figures 9a-c for Emicizumab, and in Figures 10a-f for Vanucizumab.

**Figure 9:**
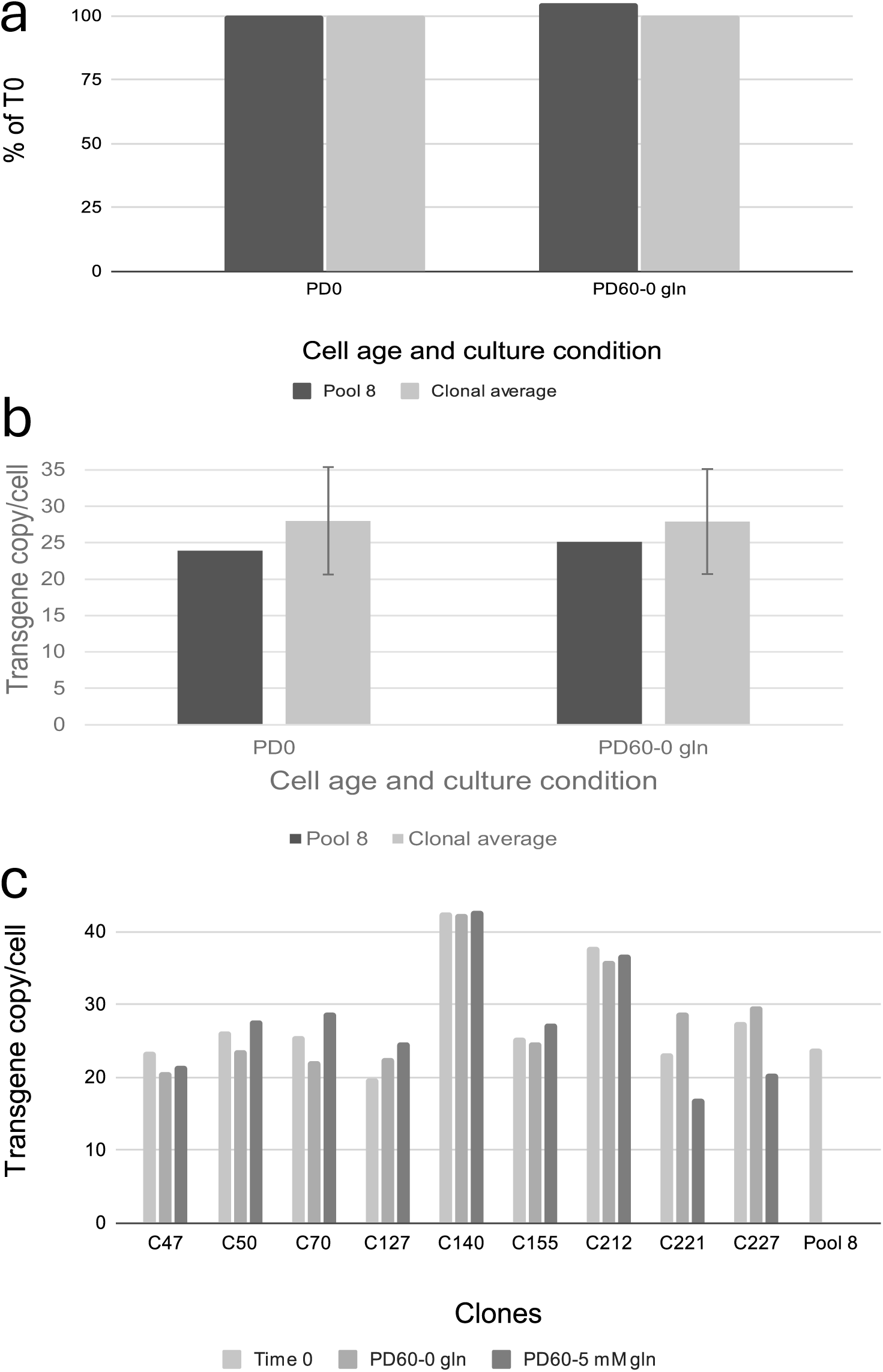
Stably integrated transgene copy-number comparability and stability in the Emicizumab expressing stable Pool 8 and its derivative clones. (a) Pool 8 and clonal average transgene copy numbers were determined at T0 and PD60. (b) PD60 Pool 8 and clonal average copy numbers expressed as % of the corresponding T0 values. (c) Clonal copy number stability and comparison of the clonal copy numbers to the T0 Pool 8 value.

**Figure 10:**
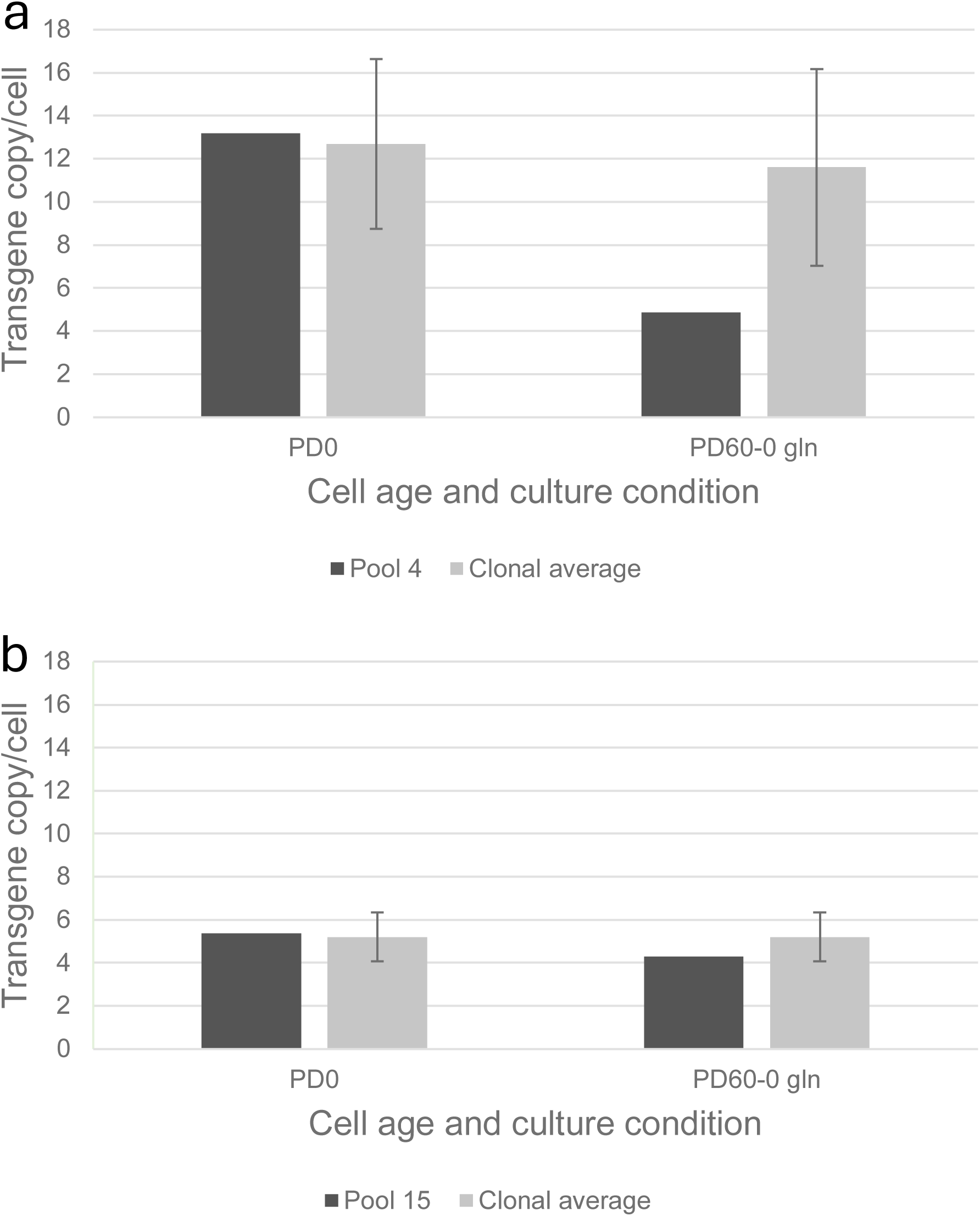
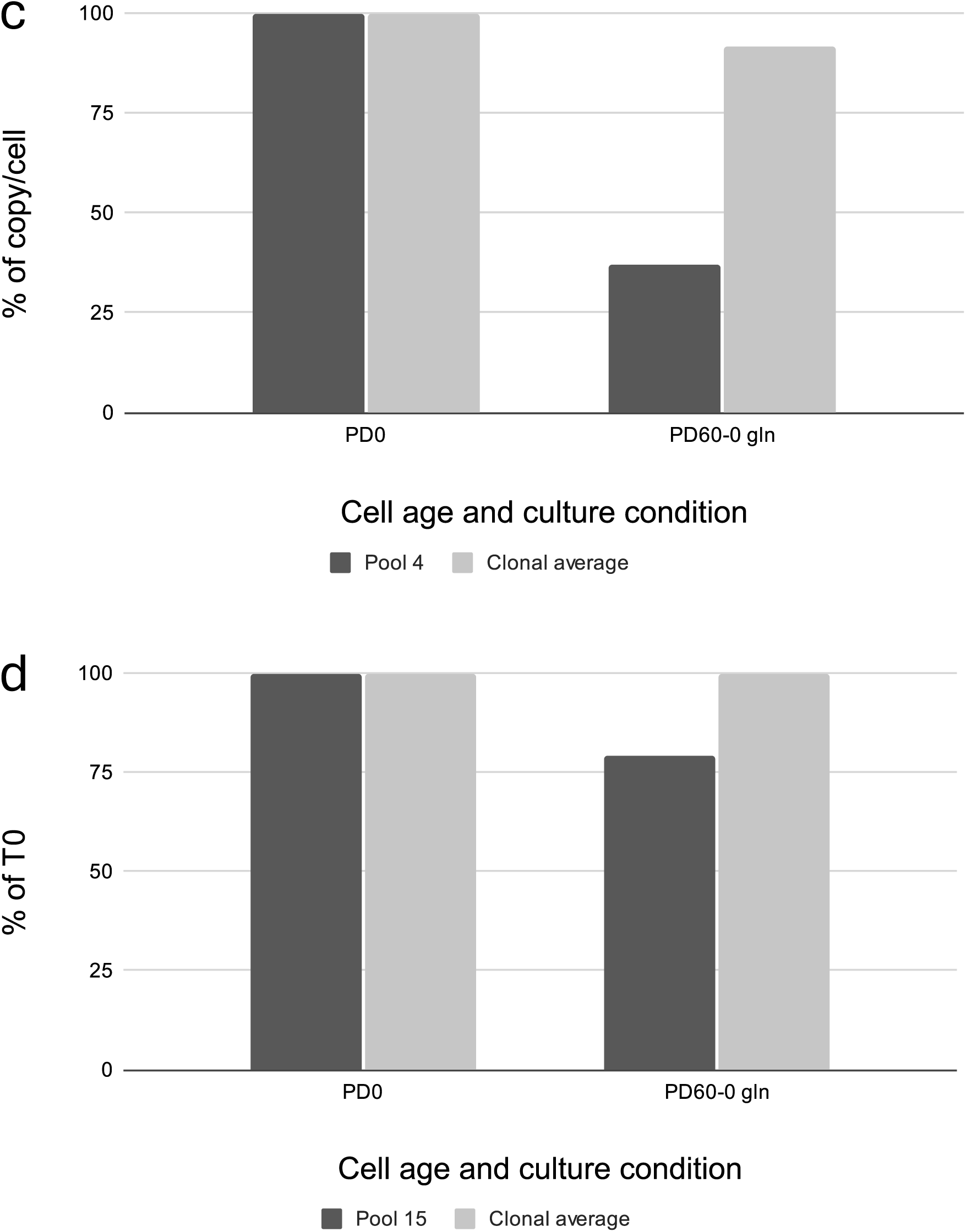
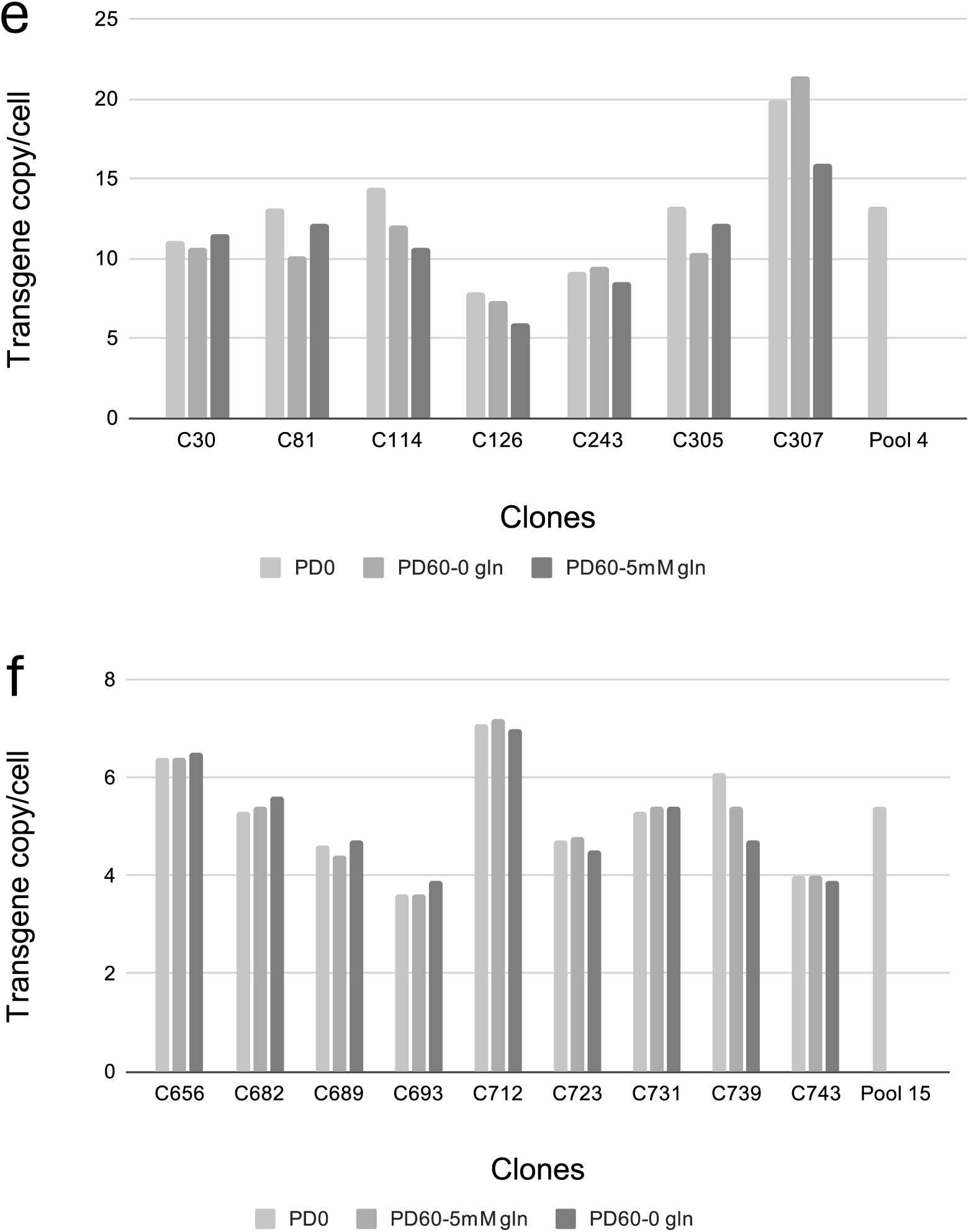
Stably integrated transgene copy-number comparability and stability in the Vanucizumab expressing stable Pools 4 and 15 and in their derivative clones. Pool 4 (a) and Pool 15 (b) and their respective clones’ average copy numbers were determined at T0 and PD60. PD60 Pool 4 (c) and Pool 15 (d) and their respective clones’ average copy-numbers expressed as % of the corresponding T0 values. Clonal copy-number stability and comparison of the clonal copy-numbers to the corresponding T0 Pool 4 (e) and Pool 15 (f) values.

The copy number differences and stability in the three pools are reviewed in the Discussion. The clonal copy numbers remain stable at least up to PD60 even without the presence of selective pressure during the stability passages.

## Discussion

The efficient, cost-effective, and rapid development of bispecific antibody-producing manufacturing cell lines remains a challenge. Since its inception, the Leap-In Transposase-mediated process has generated many bispecific antibody-producing stable pools and clones where heterodimerization is the most important CQA.

The fundamental characteristics and unique advantages of the Leap-In transposase-based cell line development platform have been described previously (Rajendran *et al*, 2021; Huhn *et al*, 2023).

Leap-In-mediated stable pools were utilized to manufacture GMP-grade diagnostic reagent antibodies (Tu, B. *et al*. 2024). An example for Leap-In transposon-based expression construct optimization, to increase productivity and heterodimer fraction, is presented in Wang, Y. *et al*, 2022.

We here demonstrate that the Leap-In transposase-mediated bispecific antibody expressing stable pools strongly predict the productivity and the heterodimer fraction expressed by the derivative clones. The almost perfect productivity stability is a hallmark of the Leap-In mediated monomer and homodimer/multimer producing clones (Rajendran, S. *et al*. 2021). The data presented herein support the extension of the previous stability statement to heterodimeric bispecific antibody-expressing clones, indicating that the selected subunit ratios are stably maintained (Figures 5c, 7e, 7f). Since the productivity of the individual stable Leap-In clones is stable and given that these clones are inherently representative of the parental stable pool, the potential productivity decrease in stable Leap-In pools is not driven by genetic instability. Instead, decreasing pool productivity is a consequence of population dynamics where faster-growing clones incrementally increase their proportion in the pool. Since faster growth correlates with lower recombinant protein production, the productivity of a growing pool will eventually decrease.

In this study, we used two model molecules: one three-chain bispecific, expressed by a 3ORF construct, and one four-chain bispecific antibody expressed using two different construct designs, a 4ORF and a 2 x 2ORF co-transfection approach. While their T0 productivity was similar, the three stable pools exhibited different productivity decrease profiles during the PD60 long stability study. The potential underlying mechanisms behind this clonal productivity distribution related population dynamics are 1) competition between cell division and recombinant protein production for cellular resources (Donaldson 2021), 2) ER stress/UPR driven cell cycle arrest (Lee 2019; Thomas 2013), and 3) UPR induced apoptosis (Prashad 2014; Chen 2023).

Emicizumab Pool 8 had the highest copy number (Figure 9a) and the best productivity stability profile (Figures 5a, 5b). Even the PD60 culture produced Emicizumab at the T0 level. This may suggest that Emicizumab overexpression is well tolerated by the cells and does not induce a significant UPR response, which in turn could adversely affect the growth of higher producer cells. This unique Leap-In pool productivity stability was associated with no copy number decrease between T0 and PD60 Pool 8 samples (Figures 9a, 9b).

The two 4-chain Vanucizumab expressing pools (Pool 4 and Pool 15), despite their copy number differences (Figures 10a, 10b), exhibited comparable T0 productivities (Figures 7a, 7b), suggesting a ceiling in the cells’ post-transcriptional processing efficiency. There was, however, a difference between the dynamics of productivity decreases in the two pools (Figures 7c, 7d). The T0 copy number of Pool 4 was higher than Pool 15 (Figures 10a, 10b), and considering that Pool 15 was established by co-transfection, the chain-specific copy number in Pool 15 is, on average, ∼half of the measured value (∼3 copies/cell). There was a higher (∼63%) copy number decrease in Pool 4, while the copies were better maintained (∼21% loss) in Pool 15 (Figures 10c, 10d). At PD60, both pools reached a similar copy number (∼4 copies/cell), indicating the lowest recombinant GS level required for survival in the glutamine-free culture condition (Figure 10b). The level of copy number decrease correlated with the corresponding productivity losses, showing a higher (∼50%) productivity loss in Pool 4 during the stability study. In comparison, the productivity decrease was ∼35% in Pool 15 by PD60 (Figs. 7c, 7d). The 3ORF Pool 8, the 4ORF Pool 4, and the 2 x 2ORF Pool 15 had similar T0 productivities. The process steps, the expression host, the expression construct elements, and the codon optimization algorithms were identical during stable pool development. This, together with the similar T0 productivities, suggests that the differences in pool productivity “stability” are driven by the molecules’ intrinsic biophysicochemical properties affecting translation, folding/assembly, and secretion.

Nevertheless, due to the highly homogeneous clonal productivity distribution demonstrated in the three pools (Figures 4a-c, Table 4), the productivity decrease in Leap-In mediated stable pools is considerably lower than those observed in pools generated by random integration. The more homogeneous clonal productivity distribution also justifies why ranking only a small number of clones, in this study 35, 47, and 55 from their respective selected pools, is sufficient to find several high producers with a satisfactory product quality profile and remarkable productivity and product quality stability (5c, 6c, 7e, 7f, 8e, 8f).

A common quality attribute for every bispecific molecular design, manufacturing cell substrate, and production process is the fraction of the desired heterodimers produced by the cells before any heterodimer enrichment step. In this study, we chose the heterodimer fraction, measured from proA capture of harvest material, as the product quality ranking attribute. The contribution of cell line development to achieve the highest unenriched heterodimer fraction is to express the various subunits at an optimal ratio, favoring heterodimer formation over the homodimer species. This optimal ratio depends on the molecular architecture, the various chain pairing choices, and the actual sequence of the subunits. Consequently, the optimal expression ratio is typically not trivial to predict. Traditional cell line development approaches rely on screening thousands of clones using high-throughput methods, sometimes with scalability limitations. In stark contrast, due to the high pool to clone comparability described in this study, characterizing a few stable pools and selecting the one with acceptable productivity and product quality ensures that the protein produced by the derivative clones will also meet expected specifications. Since the highest heterodimer fraction is driven by a delicate balance between subunit expression levels, any change in the optimal ratio will result in a decrease in the heterodimer fraction. Accordingly, the expression ratio between every individually expressed subunit should remain constant (stable) to support large-scale manufacturing. The data presented (Figs. 6a-c and 8a-f) indicate that the heterodimer fractions remained stable for at least PD60 in both the derivative clones and even in the parental stable pools. The heterodimer stability detected in the pools was independent of the pools’ productivity stability, and the heterodimer fractions remained at close to T0 level even when the pool productivity decreased. This observation is in line with the structural and functional integrity of the integrated Leap-In transposon-based expression constructs (Rajendran, S. *et al*., 2021), responsible for maintaining the subunit ratios, hence the heterodimer stability. The population dynamics-driven transgene copy number decrease in some of the pools/conditions (Figures 10a-d), while decreasing productivity, does not alter the relative individual subunit expression levels.

In summary, the Leap-In transposase-mediated stable bispecific pools’ T0 performance reliably predicts the derivative clones’ productivity and heterodimer fraction. The derivative clones demonstrated productivity stability, and the heterodimer fraction remained stable in the three model pools and clones independently of the productivity stability profile. In addition to the bispecific Leap-In pools’ predictive value, their stability profile enables manufacturing representative drug substances even at large scale. The variable productivity decrease in Leap-In mediated stable pools expressing different bispecific molecules is not associated with genetic instability, but rather depends on product sequence, design, architecture, and characteristics.

The paradigm shift in cell line development driven by the Leap-In transposase platform is here extended to heterodimeric protein architectures, where ranking a handful of pools and isolating a relatively small number (∼50) of clones from a selected pool with optimal productivity and CQA’s results in a sufficient number of clones with stable productivity and product quality profile.

## Acknowledgements

The authors would like to acknowledge members of ATUM’s Cell Line Development, Protein Purification, and Protein Analytical Departments for performing three extra CLD projects in addition to their client work. We also would like to thank Calvin Tang, Efrain Zarazua, and Jennifer Nguyen for the copy number analysis and Steve Nakazawa Hewitt for creating the illustrations.

## References

Brinkmann, U., Kontermann, R.E. (2017). The making of bispecific antibodies. mAbs. 9 (2): 182–212.

Chen, X., Shi, C., et al., (2023). Endoplasmic reticulum stress: molecular mechanisms and therapeutic targets. Signal Transduct Target Ther. 8: 352

Deshaies, R.J. (2020). Multispecific drugs herald a new era of biopharmaceutical innovation. Nature. 580 (7803): 329–338.

Donaldson, S., Dale, M.P., Rosser, S.J. (2021). Decoupling growth and protein production in CHO cells: a targeted approach. Front Bioeng Biotechnol. 9: 658325.

Godar, M., de Haard, H., et al., (2018). Therapeutic bispecific antibody formats: a patent applications review (1994-2017). Expert Opin Ther Pat. 28 (3): 251–276.

Hertel, O., Neuss, A., et al., (2022). Enhancing stability of recombinant CHO cell lines by CRISPR/Cas-9-mediated site-specific integration into regions with distinct histone modifications. Front Bioeng Biotechnol. 13 (10): 1010719.

Huhn, S.C., Chang, M., et al., (2023). Genomic features of recombinant CHO clones arising from transposon-based and randomized integration. J Biotechnol. 373: 73–81.

Kitazava, T., Shima, M. (2020). Emicizumab, a humanized bispecific antibody to coagulation factors IXa and X with a Factor VIIIa-cofactor activity. Int J Hematol. 111 (1): 20–30.

Lee, D., Hokinson, D., Park, S. (2019). ER stress induces cell cycle arrest at the G2/M phase through eIEF2α phosphorylation and GADD45α. Int J Mol Sci. 20 (24): 6393.

Madsen, A.V., Pedersen, L.E., et al., (2024). Design and engineering of bispecific antibodies: insights and practical considerations. Front Bioeng Biotechnol. 12: 1352014.

Prashad, K, Mehra, S. (2015). Dynamics of unfolded protein response in recombinant CHO cells. Cytotechnol. 67 (2): 237–254.

Rajendran, S., Balasubramanian, S., et al., (2021). Accelerating and de-risking CMC development with transposon-derived manufacturing cell lines. Biotechnol Bioeng. 118 (6): 2301–2311.

Schaefer, W., Regula, J.T., et al. (2011). Immunoglobulin domain crossover as a generic approach for the production of bispecific IgG antibodies. Proc Natl Acad Sci U S A. 108 (27): 11187–11192.

Spiess, C., Zhai, Q., Carter, P.J. (2015). Alternative molecular formats and therapeutic applications for bispecific antibodies. Mol Immunol 67: 95–106.

Thomas, S.E., Malzer, E., Ordonez, A. (2013). p53 and translational attenuation regulate distinct cell cycle checkpoints during endoplasmic reticulum (ER) stress. J Biol Chem 288 (11): 7606–7617.

Tu, B., Lin, Z., et al., (2024). Recombinant Antibody-Producing Stable CHOK1 Pool Stability Study. Monoclon Antib Immunodiagn Immunother 43 (4): 119–126.

Wang, Y., Qiu, H., et al., (2022) An innovative platform to improve asymmetric bispecific antibody assembly, purity, and expression level in stable pool and cell line development. Biochem Bioeng J. 188: 108683

